# Germinal centre-driven maturation of B cell response to SARS-CoV-2 vaccination

**DOI:** 10.1101/2021.10.31.466651

**Authors:** Wooseob Kim, Julian Q. Zhou, Alexandria J. Sturtz, Stephen C. Horvath, Aaron J. Schmitz, Tingting Lei, Elizaveta Kalaidina, Mahima Thapa, Wafaa B. Alsoussi, Alem Haile, Michael K. Klebert, Teresa Suessen, Luis Parra-Rodriguez, Philip A. Mudd, William D. Middleton, Sharlene A. Teefey, Iskra Pusic, Jane A. O’Halloran, Rachel M. Presti, Jackson S. Turner, Ali H. Ellebedy

## Abstract

Germinal centres (GC) are lymphoid structures where vaccine-responding B cells acquire affinity-enhancing somatic hypermutations (SHM), with surviving clones differentiating into memory B cells (MBCs) and long-lived bone marrow plasma cells (BMPCs)^1–4^. Induction of the latter is a hallmark of durable immunity after vaccination^5^. SARS-CoV-2 mRNA vaccination induces a robust GC response in humans^6–8^, but the maturation dynamics of GC B cells and propagation of their progeny throughout the B cell diaspora have not been elucidated. Here we show that anti-SARS-CoV-2 spike (S)-binding GC B cells were detectable in draining lymph nodes for at least six months in 10 out of 15 individuals who had received two doses of BNT162b2, a SARS-CoV-2 mRNA vaccine. Six months after vaccination, circulating S-binding MBCs were detected in all participants (n=42) and S-specific IgG-secreting BMPCs were detected in 9 out of 11 participants. Using a combined approach of single-cell RNA sequencing of responding blood and lymph node B cells from eight participants and expression of the corresponding monoclonal antibodies, we tracked the evolution of 1540 S-specific B cell clones. SHM accumulated along the B cell differentiation trajectory, with early blood plasmablasts showing the lowest frequencies, followed by MBCs and lymph node plasma cells whose SHM largely overlapped with GC B cells. By three months after vaccination, the frequency of SHM within GC B cells had doubled. Strikingly, S^+^ BMPCs detected six months after vaccination accumulated the highest level of SHM, corresponding with significantly enhanced anti-S polyclonal antibody avidity in blood at that time point. This study documents the induction of affinity-matured BMPCs after two doses of SARS-CoV-2 mRNA vaccination in humans, providing a foundation for the sustained high efficacy observed with these vaccines.

## Main text

Phase 3 trials of SARS-CoV-2 mRNA-based vaccines demonstrated the remarkable efficacy of these vaccines against coronavirus disease 2019 (COVID-19)^9,10^. The Pfizer-BioNTech SARS-CoV-2 mRNA vaccine (BNT162b2) remains highly efficacious at preventing symptomatic infection for at least six months after the initial two-dose vaccine series in individuals with no history of SARS-CoV-2 infection^11^. This is despite a decrease in S-binding antibody concentrations and neutralizing antibodies that occur during the same period^12,13^. We have previously shown that vaccination of human subjects with BNT162b2 induces a robust but transient circulating plasmablast (PB) response followed by remarkably persistent germinal centre (GC) reactions in the draining lymph nodes^6^. Whether these persistent GC responses lead to the generation of affinity-matured bone marrow-resident plasma cells remains unclear. To address this question, we undertook longitudinal follow-up of participants enrolled in our previously described observational study of 43 healthy participants (13 with a history of confirmed SARS-CoV-2 infection) who received two doses of the Pfizer-BioNTech SARS-CoV-2 mRNA vaccine (BNT162b2) (**Extended Data Tables 1, 2**)^6,7^. Long-term blood samples (n=42), fine needle aspirates (FNAs) of the draining axillary lymph nodes (n=15) and bone marrow aspirates (n=11) were collected 29 weeks after the primary vaccination (**Fig. 1a**). None of the participants who contributed FNA or bone marrow specimens had a history of SARS-CoV-2 infection.

**Figure 1.**
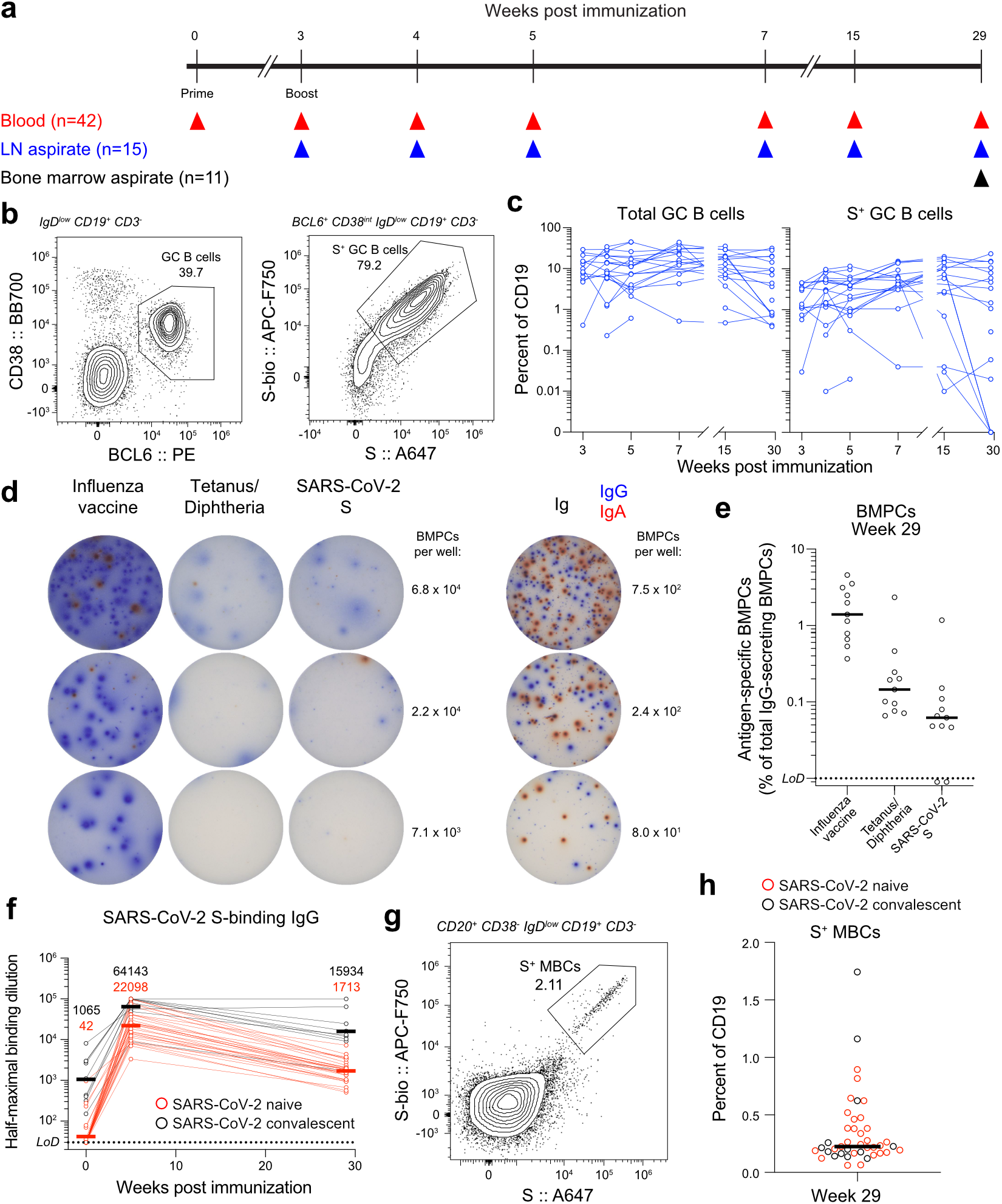
Persistence of humoral immune responses to SARS-CoV-2 mRNA vaccination. **a**, Study design. Forty-three healthy adult volunteers (13 with a history of SARS-CoV-2 infection) were enrolled, followed by BNT162b2 mRNA SARS-CoV-2 vaccination. Blood (n=42) was collected before immunization, and at 3, 4, 5, 7, 15, and 29 weeks after primary immunization. For 15 participants without a history of SARS-CoV-2 infection, aspirates of ipsilateral axillary lymph nodes were collected at 3, 4, 5, 7, 15, and 29 weeks after primary vaccination. For 11 participants without a history of SARS-CoV-2 infection, aspirates of bone marrow were collected at 29 weeks after primary vaccination. **b**, Representative flow cytometry plots of GC B cells (CD19^+^ CD3^−^ IgD^low^ BCL6^+^ CD38^int^ live singlet lymphocytes) and SARS-CoV-2 S staining on GC B cells in draining lymph nodes 29 weeks post-vaccination. **c**, Kinetics of total (left) and S^+^ GC B cells (right) as gated in **b**. **d**, Representative ELISpot wells coated with the indicated antigens or anti-immunoglobulin and developed in blue and red for IgG and IgA, respectively, after plating the indicated numbers of magnetically enriched BMPCs. **e**, Frequencies of BMPCs secreting IgG antibodies specific for the indicated antigens 29 weeks after vaccination. **f**, Plasma IgG titers against SARS-CoV-2 S measured by ELISA in participants without (red) and with (black) a history of SARS-CoV-2 infection. Horizontal lines indicate geometric means, also shown above time points. Results are from one experiment performed in duplicate. **g**, Representative flow cytometry plot of SARS-CoV-2 S staining on MBCs (CD20^+^ CD38^−^ IgD^low^ CD19^+^ CD3^−^ live singlet lymphocytes) in blood 29 weeks after primary vaccination. **h**, Frequencies of S^+^ MBCs in participants without (red) and with (black) a history of SARS-CoV-2 infection as gated in **g**. Horizontal lines indicate median values in **e** and **h**. Dotted lines indicate limits of detection in **e** and **f**. Symbols at each time point represent one sample in **c** (n=15), **e** (n=11), **f** (n=38), and **h** (n=42).

GC B cells were detected in FNAs from all 15 participants (**Fig. 1b, c, left panels, Extended Data Fig. 1a, Extended Data Table 3**). All 14 participants with FNAs collected prior to week 29 generated SARS-CoV-2 S-binding antigen-specific GC B cell responses of varying magnitudes (**Fig 1b, c, right panels, and Extended Data Table 3**). Intriguingly, S-binding GC B cells were detected in FNAs from 10 of 15 participants at week 29 (**Fig. 1b, c, right panels, Extended Data Table 3**), demonstrating that more than half of the sampled subjects maintained an antigen-specific GC B cell response more than 6 months after vaccination. S-binding LNPCs were also detected in FNAs from all 15 participants and exhibited kinetics similar to S-binding GC B cells, albeit at lower frequencies within the total B cell population (**Extended Data Fig. 1a-b, Extended Data Table 3**). None of the FNAs demonstrated significant contamination with peripheral blood based upon the nearly complete absence of CD14+ myeloid cells (**Extended Data Table 4**).

To determine whether SARS-CoV-2 mRNA vaccination induces antigen-specific bone marrow plasma cells (BMPCs), we examined bone marrow aspirates collected 29 weeks after primary vaccination. We first magnetically enriched BMPCs and then measured the frequencies of BMPCs secreting IgG or IgA antibodies against the 2019-2020 inactivated influenza virus vaccine, the tetanus-diphtheria vaccine or SARS-CoV-2 S protein by enzyme-linked immunosorbent spot assay (ELISpot) (**Fig. 1d-e, Extended Data Fig. 1c**). Influenza- and tetanus-diphtheria vaccine-specific IgG-secreting BMPCs (median frequencies of 1.4% and 0.15%, respectively) were detected in all 11 participants (**Fig. 1e**). SARS-CoV-2 S-binding IgG-secreting BMPCs were detected in 9 of 11 participants (median frequency of 0.06%). IgA-secreting BMPCs specific for the seasonal influenza vaccine were detected in 10 of 11 participants, but IgA-secreting BMPCs directed against tetanus-diphtheria-vaccine and SARS-CoV-2 S protein were largely below the limit of detection (**Extended Data Fig. 1c**). Importantly, none of the participants had detectable S-specific antibody-secreting cells (ASCs) in blood at the time of bone marrow collection, indicating that the detected BMPCs represent bona fide bone marrow-resident cells and not contamination from circulating blood PBs.

All study participants had readily detectable plasma anti-S IgG antibodies and circulating S-specific memory B cells (MBCs) at the 29 week time point (**Fig. 1f, g, and h**), consistent with the presence of S-specific BMPCs and GC B cells in the bone marrow and lymph node, respectively. Notably, anti-S IgG titers 29 weeks following vaccination were higher than titers observed in a separate cohort of unvaccinated SARS-CoV-2-infected convalescent subjects measured 29 weeks post-infection^14–16^(**Extended Data Fig. 1d, Extended Data Table 1**). Vaccinated participants with a history of confirmed SARS-CoV-2 infection showed significantly higher titers of plasma anti-S IgG 5 weeks and 29 weeks following vaccination than their naive counterparts^14,16,17^ (**Fig. 1f**). Similar trends were observed for plasma anti-S IgM and IgA (**Extended Data Fig. 1e**). SARS-CoV-2 S-binding MBCs were observed in all participants, with a median frequency of 0.23% of total CD19^+^ circulating B cells(**Fig. 1g, h, Extended Data Fig. 1f**). . Collectively, these results demonstrate that SARS-CoV-2 mRNA vaccination generates robust, persistent humoral responses.

To investigate SARS-CoV-2 S-specific B cell composition and clonal distribution within blood, lymph node and bone marrow, we performed single-cell RNA sequencing (scRNA-seq) for both transcriptomic and B cell receptor (BCR) repertoire profiling on 8 participants who contributed samples from all three of these tissue compartments. To maximize the number of blood PBs characterized by scRNA-seq, we sorted PBs from blood samples collected at the peak of the circulating PB response, 4 weeks after primary vaccination^6^(**Extended Data Fig. 2a**). We selected 7 and 15 weeks post-vaccination as representative early and late time points to interrogate immune responses in the lymph node, except that we were unable to include samples from 2 of the 8 participants at the late time points due to insufficient sample remaining. For these two subjects, we analyzed two separate relatively early time points - weeks 5 and 7 for participant 02a, and weeks 4 and 7 for participant 04. Single-cell transcriptome analysis of lymph nodes revealed distinct populations of T and B cells, NK cells, monocytes, plasmacytoid dendritic cells, and follicular dendritic cells, as previously described^18–21^(**Fig. 2a, bottom left panel, Extended Data Fig. 2b, c, Extended Data Table 5**). To further distinguish between transcriptionally distinct subsets of B cells in the lymph node following SARS-CoV-2 vaccination, we performed unbiased clustering of the B cell populations from the total cellular analysis (**Fig 2a, bottom left panel, Extended Data Fig. 2b, c**) based upon gene expression (**Fig. 2a, bottom right panel, Extended Data Fig. 2d, e, Extended Data Table 5**). Consistent with our flow cytometry results, considerable numbers of cells clustered into transcriptional groups defining GC B cells (33.8%) and LNPCs (7%) in the lymph node.

**Figure 2.**
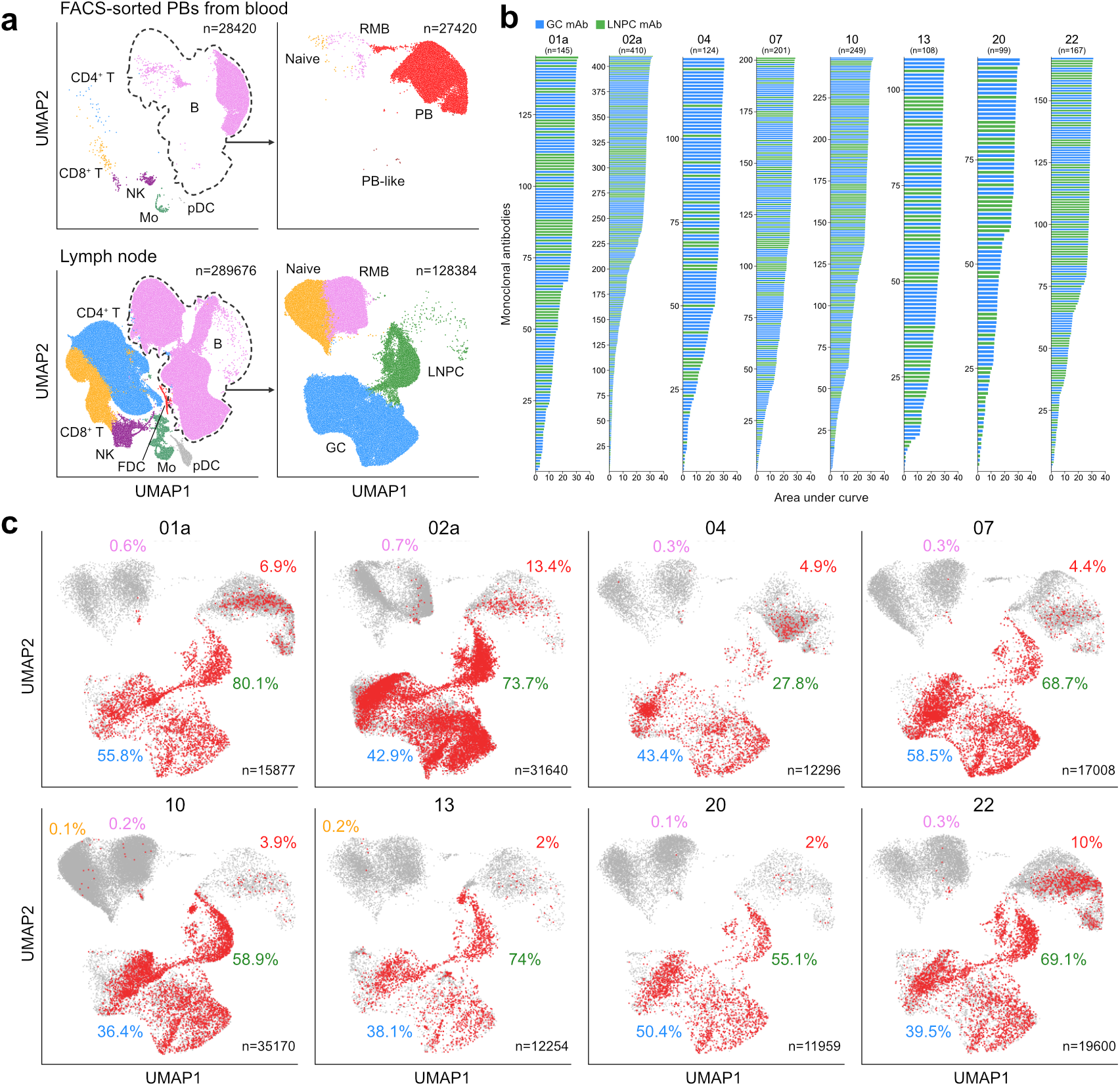
Identification of SARS-CoV-2 S-binding B cell clones in draining lymph nodes. **a**, Uniform manifold approximation and projection (UMAP) of scRNA-seq data from PBs sorted from PBMC (upper) and whole FNA of draining axillary lymph nodes (lower), and UMAP of B cell scRNA-seq clusters from each compartment (right). Each dot represents a cell, colored by phenotype as defined by the gene expression profile. Total numbers of cells are at the top right corner. FDC, follicular dendritic cell; GC, germinal center B cell; Mo, monocyte; NK, natural killer cell; LNPC, lymph node plasma cell; PB, plasmablast; pDC, plasmacytoid dendritic cell; RMB, resting memory B cell. **b**, Positive binding of recombinant monoclonal antibodies (mAbs) derived from GC B cells (blue) or LNPCs (green) to SARS-CoV-2 S measured by ELISA. Areas under the curve were calculated by setting the mean + three times the s.d. of background binding to bovine serum albumin (BSA) as a baseline. Results are from one experiment performed in duplicate. **c**, SARS-CoV-2 S-binding clones visualized on UMAP of B cell clusters in each participant. Percentages are of SARS-CoV-2 S-binding clones within GC B cells (blue), LNPCs (green), PBs (red), RMBs (pink) or naive B cells (yellow). Total numbers of cells are at the bottom right corner.

To achieve reasonably comprehensive estimation of the vaccine-responding B cell clones in the lymph nodes, we selected one representative BCR from either the GC B cell or the LNPC transcriptional clusters for recombinant monoclonal antibody (mAb) generation from every single-cell sequenced BCR clone containing at least 3 cells and present in one or both compartments. A total of 2099 recombinant mAb from 8 participants were generated and subsequently tested for binding to SARS-CoV-2 S by ELISA. We found that 1503 of the 2099 recombinant mAbs bound to SARS-CoV-2 S (**Fig. 2b, Extended Data Table 6**). In subsequent analyses of these S-binding BCRs, we also included 15, 5 and 17 previously identified and published S-binding mAbs generated from GC B cells at week 4 from, respectively, participants 07, 20, and 22^6^. Clonal relationships among S-specific BCRs were analyzed using heavy chains from scRNA-seq libraries (**Extended Data Table 7**), bulk-seq libraries for sorted GC B cells and LNPCs (**Extended Data Fig. 2f, Extended Data Table 8**), as well as previously published bulk-seq libraries of sorted PBs and GC B cells^6^ and magnetically enriched IgD^low^ activated or memory B cells from PBMC^22^. B cell clones consisting of experimentally-validated S-binding B cells accounted for 44.5% and 67.3%, respectively of the single-cell profiled GC B cells and LNPCs (**Fig. 2c, Extended Data Fig. 2g, Extended Data Table 5**). B cells that were clonally related to S-binding B cells were also found in the PB cluster in blood (6.7%) and resting memory B cell (RMB) cluster in lymph nodes (0.3%) (**Fig. 2c, Extended Data Fig. 2g, Extended Data Table 5**). Thus, our results clearly demonstrate robust induction of SARS-CoV-2 S-binding GC B cells and LNPCs in the lymph node at the single-cell level following mRNA vaccination.

We next analyzed the proportion of S-binding GC B cells clonally related to PBs in the blood at week 4. The frequencies of PB-related S-binding GC B cells varied broadly among participants and over time from 11.5% to 82.5% (**Fig. 3a**). BCR sequences of S-binding GC B cells had significantly higher levels of somatic hypermutation (SHM) compared to clonally related PB, and this difference increased over time (**Fig. 3b**). We observed a 2.5-fold increase in SHM frequency among all S-binding GC B cells between weeks 4 and 15 (**Fig. 3c, Extended Data Fig. 3a**). We also observed that the relative proportion of IgA-expressing S-binding GC B cells increased in the lymph node over time (**Fig. 3d**). Collectively, these results suggest that GC B cells continue to evolve for at least 15 weeks following mRNA vaccination.

**Figure 3.**
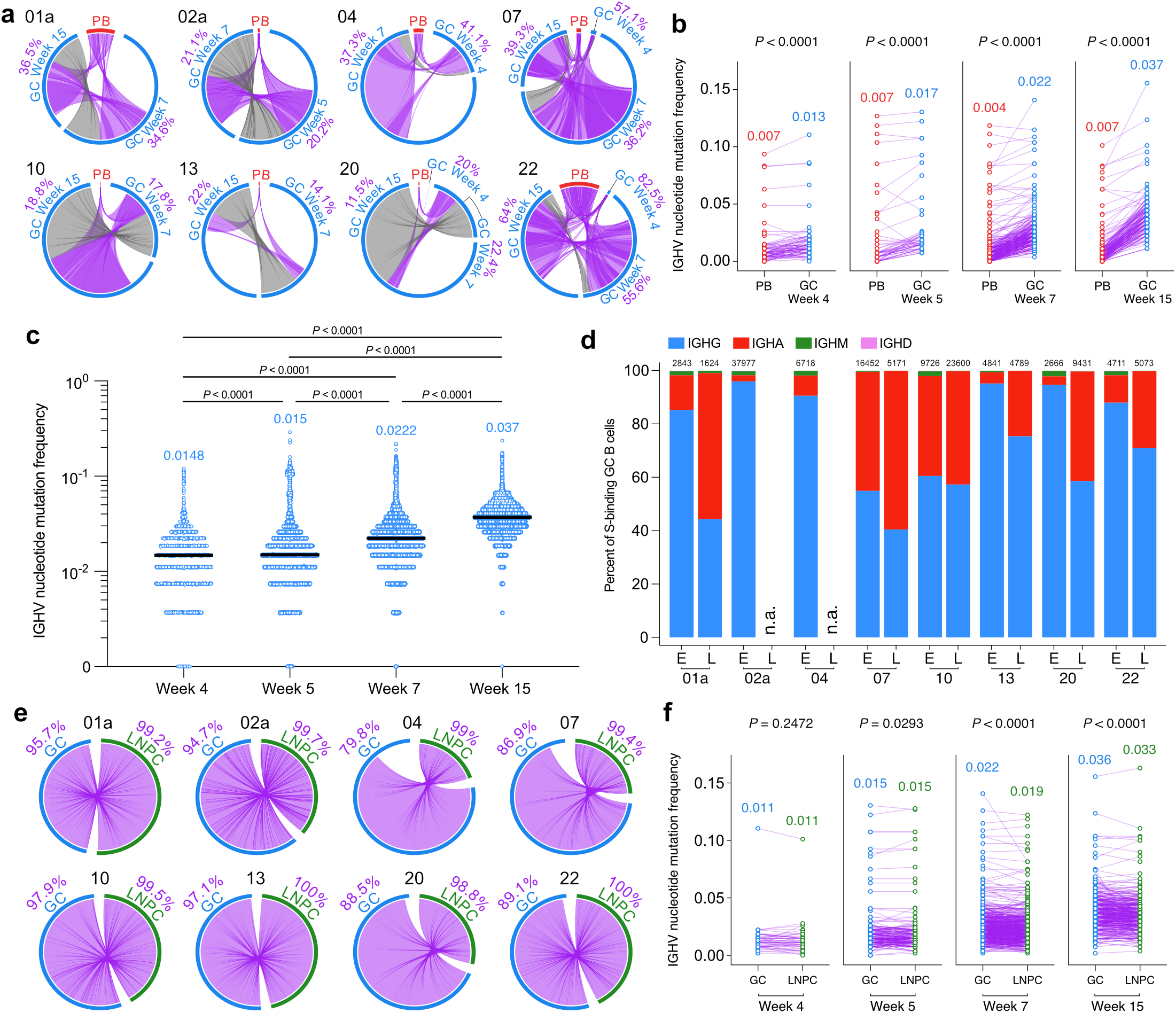
Maturation of SARS-CoV-2 S-binding B cells in the lymph node. **a**, Clonal overlap of SARS-CoV-2 S-binding sequences from bulk and single-cell BCR repertoire analysis between PBs sorted from blood 4 week post-vaccination and GC B cells at indicated time points. Arc length corresponds to the number of BCR sequences and chord width corresponds to clonal group size. Purple chords correspond to clones spanning both the PB and GC compartments. Grey chords correspond to clones spanning only the GC compartment. Percentages are of GC B cell clones related to PBs at each time point. **b**, Comparison of immunoglobulin heavy chain variable (*IGHV*) region nucleotide mutation frequency of clonally related, SARS-CoV-2 S-binding PBs and GC B cell clones that are clonally linked at the indicated time points. Median values are presented on the top of each data set. **c**, Comparison of *IGHV* nucleotide mutation frequency of SARS-CoV-2 S-binding GC B cells at the indicated time points. Horizontal lines and colored numbers represent median values. *P* values were determined by Kruskal-Wallis test followed by Dunn’s multiple comparison test. **d**, Percentages of GC B cells expressing IGHG (blue), IGHA (red), IGHM (green) or IGHD (pink) at the early (E) or the late (L) time point. The early time point represents 4, 5 or 7 weeks after immunization. The late time point represents 15 weeks post-immunization. Total cell numbers are on the top of each column. **e**, Circos diagram showing overlap of S-binding clonal groups between GC B cells and LNPCs over combined time points. Arc length corresponds to the number of BCR sequences and chord width corresponds to clonal group size. Purple chords link GC B cells to clonally overlapping LNPCs. Percentages are of GC B cell clones overlapping with LNPCs or *vice versa* in each participant. **f**, Comparison of *IGHV* nucleotide mutation frequency of clonally related, S-binding GC B cell and LNPCat the indicated time points. Median values are presented on the top of each data set. Each dot represents the median SHM of a clonal group within the indicated compartment in **b** and **f**. *P* values were determined by paired two-sided non-parametric Mann-Whitney test corrected for multiple testing using Benjamini and Hochberg’s method in **b** and **f**.

Clonal analysis revealed a very high degree of overlap between S-binding GC and LNPC compartments (**Fig. 3e**). Furthermore, SHM frequencies of both S-binding LNPCs and GC B cells increased over time at a remarkably similar rate with small differences in contrast to those between S-binding PB and GC B cells (**Fig. 3f**). Taken together, the near-complete clonal overlap of S-binding LNPCs with GC B cells and their lockstep accumulation of SHM suggests that LNPCs differentiate from GC B cells.

To determine whether the increase in SHM frequencies of S-specific GC B cells and LNPCs over time is reflected in increased circulating anti-S antibody binding affinity, we measured the avidity of plasma anti-S IgG at 5 and 29 weeks post-vaccination. In participants without a history of SARS-CoV-2 infection, avidity of plasma anti-S IgG increased at 29 weeks compared to 5 weeks post-vaccination. Interestingly, participants with a history of SARS-CoV-2 infection had comparable plasma anti-S IgG avidity at 5 and 29 weeks post-vaccination (**Fig. 4a**). As expected, S-binding ASCs sampled at various time-points post-vaccination exhibited higher frequencies of SHM over time, and BMPCs sampled 29 weeks post-vaccination exhibited the highest degree of SHM (**Fig. 4b, Extended Data Fig. 4a**). To understand the evolutionary relationships between S-specific ASC lineages, we analyzed S-binding clones using a hierarchical phylogenetic model. The resulting phylogenetic trees revealed a close relationship between BMPCs and LNPCs. The closely-related BMPCs and LNPCs shared common PB ancestors (**Fig. 4c**). Together these results provide strong support for a model where SARS-CoV-2 S-specific BMPCs are the products of affinity-matured GCs.

**Figure 4.**
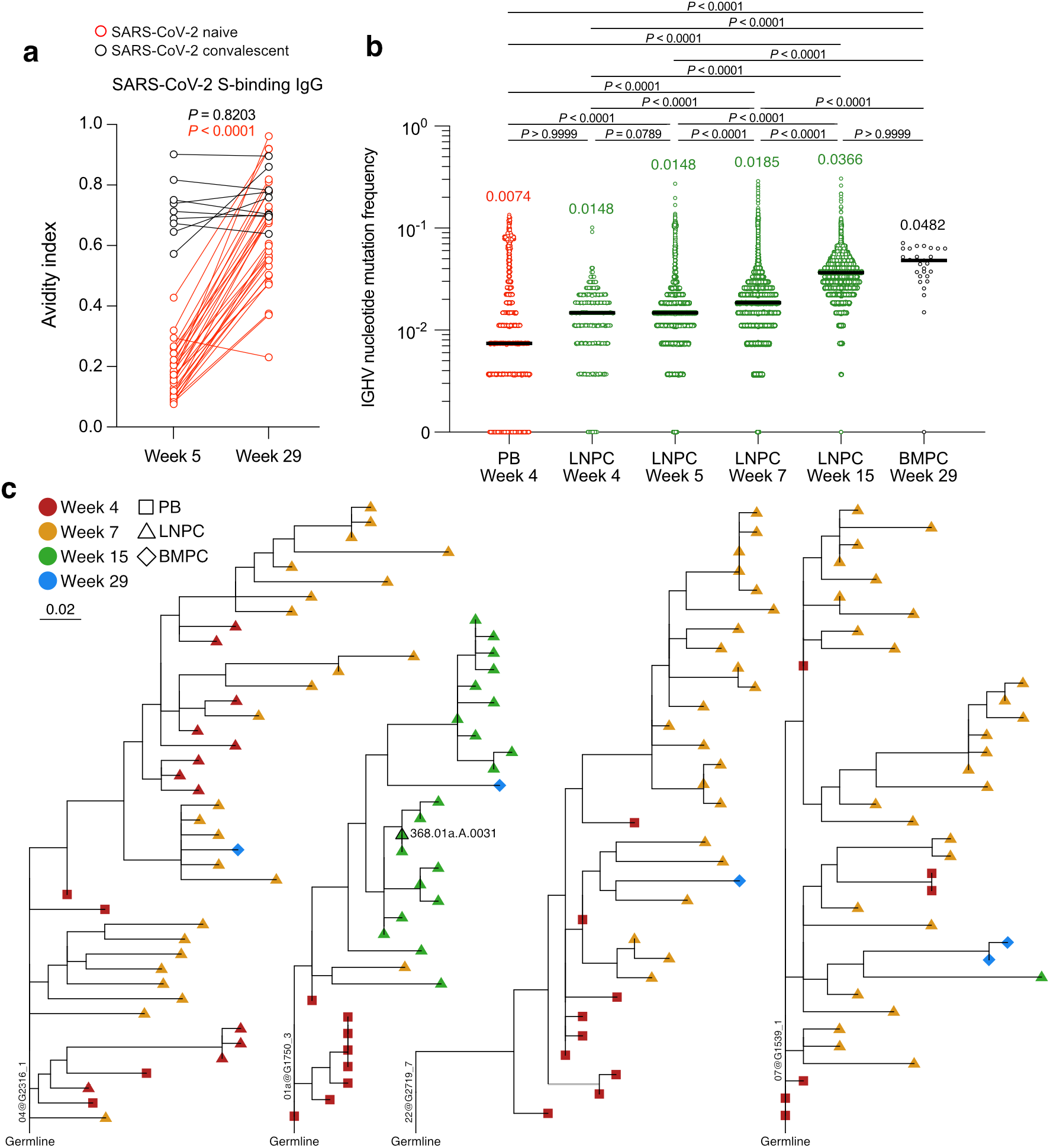
Evolution of B cell clones induced by SARS-CoV-2 vaccination. **a**, Comparison of avidity indices of plasma IgG against SARS-CoV-2 S between the indicated time points in participants without (red) and with (black) a history of SARS-CoV-2 infection. Avidity index was defined as the ratio of optical density values obtained in the presence and absence of 8M urea. Data are *P* values were determined by Wilcoxon matched-pairs signed rank test. Results are from one experiment performed in duplicate. **b**, *IGHV* nucleotide mutation frequency of S-binding PB, LNPC, and BMPC at the indicated time points. Horizontal lines and colored numbers represent median values. *P* values were determined by Kruskal-Wallis test followed by Dunn’s multiple comparison test. **c**, Phylogenetic trees of each clonal group showing inferred relations between PB (squares), LNPC (triangles), and BMPC (diamonds). ELISA-verified S-binding mAb IDs and clonal group IDs are indicated in the diagram. Horizontal branch length represents the expected number of substitutions per codon in V-region genes, corresponding to the key.

## Discussion

Upon antigenic stimulation, naïve B cells and pre-existing MBCs can differentiate into short-lived PBs that secrete relatively low-affinity antibodies or into GC B cells where they accumulate SHM and undergo affinity-based selection, with surviving clones differentiating into MBCs and long-lived BMPCs^1–3,18,23,24^. This study evaluated whether the persistent GC response induced by SARS-CoV-2 mRNA-based vaccines in humans^6^ results in generation of affinity-matured antigen-specific BMPCs. The two-dose series of BNT162b2 induced a robust SARS-CoV-2 S-binding GC B cell response that was detected in 10 of the 15 participants’ draining lymph nodes for at least six months after vaccination. The fruits of such persistent GC reactions were evident in the form of circulating SARS-CoV-2 S^+^ MBCs in all participants and SARS-CoV-2 S^+^ BMPCs detected in bone marrow specimens from 9 of the 11 participants examined six months after vaccination. It is likely that S^+^ BMPCs in those two participants are present at very low frequencies below the assay detection limit. Anti-S IgG plasma antibodies declined from an initial peak 2 weeks after secondary vaccination but remained readily detectable for at least six months after vaccination. Longitudinal tracking of over 1500 vaccine-induced B cell clones revealed the gradual accumulation of SHM within the GC B cell compartment over time. Additionally, it revealed the differentiation of those GC B cells into LNPCs within the lymph node, and then into affinity matured BMPCs. The maturity of BMPCs was reflected in the significantly enhanced avidity of circulating anti-S IgG antibodies detected at six months compared to that measured at 2 weeks after the second vaccination. Our data corroborate multiple reports demonstrating the maturation of B cell responses after SARS-CoV-2 mRNA vaccination in humans^14,15,17,25–27^. A potential limitation to our analyses of S-binding clones is that our selection strategy may have excluded some low-abundance or low-affinity S-specific clones. Nonetheless, we were able to account for 45% and 67% of all GC B cell and LNPC clones, respectively identified by scRNA-seq.

This is the first study to provide direct evidence for the induction of antigen-specific BMPCs by an mRNA-based vaccine in humans. Notably, none of the 11 participants from whom post-vaccination bone marrow specimens were examined had a history of SARS-CoV-2 infection. BMPCs that recognized contemporary seasonal influenza virus vaccine antigens and diphtheria/tetanus vaccine antigens were present at frequencies roughly 10- and 2-fold greater than those against SARS-CoV-2 S, respectively. This is likely due to both the greater number of antigenic targets contained in the former vaccines and the repeated exposures to influenza and tetanus/diphtheria vaccine antigens our study participants likely experienced in comparison to the initial exposure to the novel SARS-CoV-2 S antigen. There are some epitopes within the S protein that are conserved between human seasonal coronaviruses and SARS-CoV-2^28,29^. Cross-reactive B cells targeting these epitopes participate in PB and GC B cell responses to SARS-CoV-2 vaccination^6,30^. It is unlikely, however, that a substantial proportion of the SARS-CoV-2 S^+^ BMPCs we observed six months after immunization were part of a pre-existing pool of BMPCs, as in a previous study we could not detect any SARS-CoV-2 S^+^ BMPCs in specimens from 11 healthy individuals with no history of SARS-CoV-2 infection or vaccination^24^. It is yet to be determined, however, whether vaccine-induced SARS-CoV-2 S^+^ BMPCs we detected are indeed long-lived.

An intriguing finding in our study is that the SARS-CoV-2 S^+^ BMPCs detected six months after vaccination exhibited high SHM frequencies relative to the other B cell compartments. These data corroborate similar observations made in the mouse model^31,32^. The murine data led to a proposal of a division of labor between memory B cells and long-lived BMPCs^33,34^. Under that framework, BMPCs secrete highly specific, high-affinity antibodies that provide the first layer of protection against the invading pathogen while MBCs would only be engaged in the event that a pathogen is not fully neutralized by BMPC-derived antibodies. Consistent with this notion, multiple reports have recently documented the evolution of circulating MBCs induced by SARS-CoV-2 mRNA vaccination in humans^14,15,17,26^. These reports have shown that not only the frequency of circulating S^+^ MBCs increased over time, but their ability to recognize S proteins from emerging SARS-CoV-2 variants seems to have expanded as well^25,26^. These data indicate an important role for affinity maturation of responding B cell clones beyond increasing binding affinity to the immunizing antigen. It is yet to be elucidated whether a similar division of labor is established in human B cell responses to rapidly changing pathogens.

Our study raises a number of important questions that will need to be addressed in future studies concerning the effects of an additional homologous or heterologous immunization on the dynamics and products of ongoing germinal centres, particularly with respect to affinity and specificity to original SARS-CoV-2 S and variant strains. Overall, our data demonstrate the remarkable capacity of SARS-CoV-2 mRNA-based vaccines to induce robust and persistent GC reactions that culminate in highly matured BMPC populations. mRNA-based vaccines represent a relatively new platform that can be leveraged to rationally design new vaccines against historically challenging targets, such as influenza, RSV, HIV, and malaria. Our studies provide a foundation for how such vaccines can enhance the affinity and durability of elicited antibody responses against these challenging human pathogens.

## Materials and Methods

### Sample collection, preparation, and storage

All studies were approved by the Institutional Review Board of Washington University in St Louis. Written consent was obtained from all participants. Forty-three healthy volunteers were enrolled, of whom 13 had a history of confirmed SARS-CoV-2 infection (Extended Data Table 1, 2). Fifteen out of 43 healthy participants provided FNAs of draining axillary lymph nodes. In 6 out of the 15 participants, a second draining lymph node was identified and sampled following secondary immunization. One participant (15) received the boost vaccination in the contralateral arm; draining lymph nodes were identified and sampled on both sides. Eleven out of 43 healthy participants provided bone marrow aspirates. Forty-eight participants who had recovered from mild SARS-CoV-2 infection but had not been vaccinated within 7 months of illness were previously described^24^ (Extended Data Table 1).

Peripheral blood samples were collected in EDTA tubes, and PBMCs were enriched by density gradient centrifugation over Ficoll-Paque PLUS (Cytiva) or Lymphopure (BioLegend). The residual red blood cells were lysed with ammonium chloride lysis buffer, and cells were immediately used or cryopreserved in 10% dimethyl sulfoxide in fetal bovine serum (FBS).

Ultrasound-guided FNA of draining axillary lymph nodes was performed by a radiologist or a qualified physician’s assistant under the supervision of a radiologist. Scans were performed with a commercially available ultrasound unit (Loqic E10, General Electric, Milwaukee, WI) using an L2-9 linear array transducer with transmit frequencies of 7, 8, and 9 MHz or a L6-15 linear array transducer with transmit frequencies of 10, 12, and 15 MHz. Lymph node dimensions and cortical thickness were measured, and the presence and degree of cortical vascularity and location of the lymph node relative to the axillary vein were determined before each FNA. For each FNA sample, six passes were made under continuous real-time ultrasound guidance using 25-gauge needles, each of which was flushed with 3 ml of RPMI 1640 supplemented with 10% FBS and 100 U/ml penicillin-streptomycin, followed by three 1 ml rinses. Red blood cells were lysed with ammonium chloride buffer (Lonza), washed with washing buffer (phosphate-buffered saline supplemented with 2% FBS and 2 mM EDTA), and immediately used or cryopreserved in 10% dimethyl sulfoxide in FBS. Participants reported no adverse effects from phlebotomies or serial FNAs.

Bone marrow aspirates of approximately 30ml were collected in EDTA tubes from the iliac crest. Bone marrow mononuclear cells (BMMCs) were enriched by density gradient centrifugation over Ficoll-Paque PLUS, and then the remaining red blood cells were lysed with ammonium chloride buffer (Lonza) and washed with washing buffer. BMPCs were enriched from bone marrow mononuclear cells using EasySep Human CD138 Positive Selection Kit II (StemCell Technologies) and immediately used for ELISpot or cryopreserved in 10% dimethyl sulfoxide in FBS.

### Antigens

Recombinant soluble spike (S) protein derived from SARS-CoV-2 was expressed as previously described^35^. In brief, a mammalian cell codon-optimized nucleotide sequences coding for the soluble version of S (GenBank: MN908947.3, amino acids 1-1,213) including a C-terminal thrombin cleavage site, T4 fold trimerization domain and hexahistidine tag was cloned into the mammalian expression vector pCAGGS. The S protein sequence was modified to remove the polybasic cleavage site (RRAR to A) and two stabilizing mutations were introduced (K986P and V987P, wild-type numbering). Recombinant proteins were produced in Expi293F cells (Thermo Fisher Scientific) by transfection with purified plasmid using the ExpiFectamine 293 Transfection Kit (Thermo Fisher Scientific). Supernatants from transfected cells were collected 3 days after transfection, and recombinant proteins were purified using Ni-NTA agarose (Thermo Fisher Scientific), then buffer-exchanged into PBS and concentrated using Amicon Ultra centrifugal filters (MilliporeSigma). For flow cytometry staining, recombinant S was labeled with Alexa Fluor 7647-NHS ester or biotinylated using the EZ-Link Micro NHS-PEG4-Biotinylation Kit (Thermo Fisher Scientific); excess Alexa Fluor 647 and biotin were removed using 7-kDa Zeba desalting columns (Thermo Fisher Scientific).

### Flow cytometry and cell sorting

Staining for flow cytometry analysis and sorting was performed using freshly isolated or cryo-preserved PBMCs or FNAs. For FNA staining, cells were incubated for 30 min on ice with biotinylated and Alexa Fluor 647-conjugated recombinant soluble S and PD-1-BB515 (EH12.1, BD Horizon, 1:100) in 2% FBS and 2 mM EDTA in PBS (P2), washed twice, then stained for 30 min on ice with IgG-BV480 (goat polyclonal, Jackson ImmunoResearch, 1:100), IgA-FITC (M24A, Millipore, 1:500), CD45-A532 (HI30, Thermo, 1:50), CD38-BB700 (HIT2, BD Horizon, 1:500), CD20-Pacific Blue (2H7, 1:400), CD27-BV510 (O323, 1:50), CD8-BV570 (RPA-T8, 1:200), IgM-BV605 (MHM-88, 1:100), HLA-DR-BV650 (L243, 1:100), CD19-BV750 (HIB19, 1:100), CXCR5-PE-Dazzle 594 (J252D4, 1:50), IgD-PE-Cy5 (IA6-2, 1:200), CD14-PerCP (HCD14, 1:50), CD71-PE-Cy7 (CY1G4, 1:400), CD4-Spark685 (SK3, 1:200), streptavidin-APC-Fire750, CD3-APC-Fire810 (SK7, 1:50) and Zombie NIR (all BioLegend) diluted in Brilliant Staining buffer (BD Horizon). Cells were washed twice with P2, fixed for 1 h at 25 °C using the True Nuclear fixation kit (BioLegend), washed twice with True Nuclear Permeabilization/Wash buffer, stained with FOXP3-BV421 (206D, BioLegend, 1:15), Ki-67-BV711 (Ki-67, BioLegend, 1:200), T-bet-BV785 (4B10, BioLegend, 1:400), BCL6-PE (K112-91, BD Pharmingen, 1:25), and BLIMP1-A700 (646702, R&D, 1:50) for 1 h at 25 °C, washed twice with True Nuclear Permeabilization/Wash buffer and resuspended in P2 for acquisition. For memory B cell staining, PBMC were incubated for 30 min on ice with biotinylated and Alexa Fluor 647-conjugated recombinant soluble S in P2, washed twice, then stained for 30 min on ice with IgG-BV480 (goat polyclonal, Jackson ImmunoResearch, 1:100), IgD-Super Bright 702 (IA6-2, Thermo, 1:50), IgA-FITC (M24A, Millipore, 1:500), CD45-A532 (HI30, Thermo, 1:50), CD38-BB700 (HIT2, BD Horizon, 1:500), CD24-BV421 (ML5, 1:100), CD20-Pacific Blue (2H7, 1:400), CD27-BV510 (O323, 1:50), CD8-BV570 (RPA-T8, 1:200), IgM-BV605 (MHM-88, 1:100), CD19-BV750 (HIB19, 1:100), FcRL5-PE (509f6, 1:100), CXCR5-PE-Dazzle 594 (J252D4, 1:50), CD14-PerCP (HCD14, 1:50), CD71-PE-Cy7 (CY1G4, 1:400), CD4-Spark685 (SK3, 1:200), streptavidin-APC-Fire750, CD3-APC-Fire810 (SK7, 1:50) and Zombie NIR (all BioLegend) diluted in Brilliant Staining buffer (BD Horizon). Cells were washed twice with P2 and resuspended in P2 for acquisition. All samples were acquired on an Aurora using SpectroFlo v.2.2 (Cytek). Flow cytometry data were analyzed using FlowJo v.10 (BD Biosciences).

For sorting PBs from peripheral blood, B cells were enriched from PBMC by first using EasySep Human Pan-B cell Enrichment Kit (StemCell Technologies), and then stained with CD20-PB (2H7, 1:400), CD3-FITC (HIT3a, 1:200), IgD-PerCP-Cy5.5 (IA6-2, 1:200), CD71-PE (CY1G4, 1:400), CD38-PE-Cy7 (HIT2, 1:200), CD19-APC (HIB19, 1:200) and Zombie Aqua (all BioLegend). For sorting GC B cells and LNPCs from the lymph node, single-cell suspensions were stained for 30min on ice with PD-1-BB515 (EH12.1, BD Horizon, 1:100), CD20-Pacific Blue (2H7, 1:100), IgD-PerCP-Cy5.5 (IA6-2, 1:200), CD19-PE (HIB19, 1:200), CXCR5-PE-Dazzle 594 (J252D4, 1:50), CD38-PE-Cy7 (HIT2, 1:200), CD4-Alexa-Fluor-700 (SK3, 1:400), CD71-APC (CY1G4, 1:100), and Zombie Aqua (all BioLegend). Cells were washed twice, and single PBs (live singlet CD19^+^ CD4^−^ IgD^low^ CD38^+^ CD20^−^ CD71^+^), GC B cells (live singlet CD19^+^ CD4^−^ IgD^low^ CD71^+^ CD38^int^ CD20^+^ CXCR5^+^), LNPCs (live singlet CD19^+^ CD4^−^ IgD^low^ CD38^+^ CD20^−^CD71^+^) were sorted using a FACSAria II.

### ELISA

Assays were performed in MaxiSorp 96-well plates (Thermo Fisher) coated with 100 ul of recombinant SARS-CoV-2 S, Donkey anti-human IgG (H+L) antibody (Jackson ImmunoResearch, 709-005-149) or bovine serum albumin diluted to 1ug/ml in PBS, and plates were incubated at 4 ℃ overnight. Plates then were blocked with 10% FBS and 0.05% Tween 20 in PBS. Plasma or purified monoclonal antibodies were serially diluted in blocking buffer and added to the plates. Monoclonal antibodies and plasma samples were tested at 10 ug/ml and 1:30 starting dilution, respectively, followed by 7 additional 3-fold serial dilutions. Plates were incubated for 90 min at room temperature and then washed 3 times with 0.05% Tween 20 in PBS. Secondary antibodies were diluted in blocking buffer before adding to wells and incubating for 60 min at room temperature. HRP-conjugated goat anti-human IgG (H+L) antibody (Jackson ImmunoResearch, 109-035-088, 1:2500) was used to detect monoclonal antibodies. HRP-conjugated goat anti-Human IgG Fcγ fragment (Jackson ImmunoResearch, 109-035-190, 1:1500), HRP-conjugated goat anti-human serum IgA α chain (Jackson ImmunoResearch, 109-035-011, 1:2500), and HRP-conjugated goat anti-human IgM (Caltag, H15007, 1:4000) were used to detect plasma antibodies. Plates were washed 3 times with PBST and 3 times with PBS before the addition of *o*-phenylenediamine dihydrochloride peroxidase substrate (MilliporeSigma). Reactions were stopped by the addition of 1M hydrochloric acid. Optical density measurements were taken at 490 nm. The threshold of positivity for recombinant mAbs was set as two times the optical density of background binding to BSA at the highest concentration of each mAb. The area under the curve for each monoclonal antibody and half-maximal binding dilution for each plasma sample were calculated using GraphPad Prism v.9. Plasma antibody avidity was measured as previously described^36^. Briefly, plasma dilutions that would give an optical density reading of 2.5 were calculated from the serial dilution ELISA. S-coated plates were incubated with this plasma dilution as above and then washed one time for 5 minutes with either PBS or 8M urea in PBS, followed by 3 washes with PBST and developed as above. The avidity index was calculated for each sample as the optical density ratio of the urea-washed to PBS-washed wells.

### ELISpot

ELISpot plates were coated overnight at 4 ℃ with Flucelvax Quadrivalent 2019/2020 seasonal influenza virus vaccine (Seqirus), tetanus/diphtheria vaccine (Grifols), SARS-CoV-2 S, anti-human Ig (Cellular Technology Limited) and bovine serum albumin. A direct ex vivo ELISpot assay was performed to determine the number of total, vaccine-binding or recombinant S-binding IgG- and IgA-secreting cells present in PBMCs or enriched BMPCs using Human IgA/IgG double-color ELISpot kits (Cellular Technology Limited) according to the manufacturer’s protocol. ELISpot plates were analyzed using an ELISpot analyzer (Cellular Technology Limited).

### Single-cell RNA-seq library preparation and sequencing

Sorted PBs and whole FNA from each time point were processed using the following 10x Genomics kits: Chromium Next GEM Single Cell 5’ Kit v2 (PN-1000263); Chromium Next GEM Chip K Single Cell Kit (PN-1000286); BCR Amplification Kit (PN-1000253); Dual Index Kit TT Set A (PN-1000215). Chromium Single Cell 5’ Gene Expression Dual Index libraries and Chromium Single Cell V(D)J Dual Index libraries were prepared according to manufacturer’s instructions without modifications. Both gene expression and V(D)J libraries were sequenced on a Novaseq S4 (Illumina), targeting a median sequencing depth of 50,000 and 5,000 read pairs per cell, respectively.

### Bulk B cell receptor sequencing

RNA was prepared from sorted GC B cells and LNPCs from FNA or enriched BMPCs from bone marrow using the RNeasy Plus Micro kit (Qiagen). Libraries were prepared using the NEBNext Immune Sequencing Kit for Human (New England Biolabs) according to the manufacturer’s instructions without modifications. High-throughput 2×300-bp paired-end sequencing was performed on the Illumina MiSeq platform with a 30% PhiX spike-in according to manufacturer’s recommendations, except for performing 325 cycles for read 1 and 275 cycles for read 2.

### Preprocessing of bulk sequencing BCR reads

Preprocessing of demultiplexed pair-end reads were performed using pRESTO v.0.6.2^37^ as previously described^6^, with the exception that sequencing errors were corrected using the UMIs as they were without additional clustering (Extended Data Table 8). Previously preprocessed unique consensus sequences from reported samples^6^ were included as they were. Previously preprocessed unique consensus sequences from reported samples^22^ were subset to those with at least two contributing reads and included.

### Preprocessing of 10× Genomics single-cell BCR reads

Demultiplexed pair-end FASTQ reads were preprocessed using the ‘cellranger vdj’ command from 10× Genomics’ Cell Ranger v.6.0.1 for alignment against the GRCh38 human reference v.5.0.0 (‘refdata-cellranger-vdj-GRCh38-alts-ensembl-5.0.0’). The resultant ‘filtered_contig.fasta’ files were used as preprocessed single-cell BCR reads (Extended Data Table 7).

### V(D)J gene annotation and genotyping

Initial germline V(D)J gene annotation was performed on the preprocessed BCRs using IgBLAST v.1.17.1^38^ with IMGT/GENE-DB release 202113-2^39^. IgBLAST output was parsed using Change-O v.1.0.2^40^. For the single-cell BCRs, isotype annotation was pulled from the ‘c_call’ column in the ‘filtered_contig_annotations.csv’ files outputted by Cell Ranger.

For both bulk and single-cell BCRs, sequence-level quality control was performed, requiring each sequence to have non-empty V and J gene annotations; exhibit chain consistency in all annotations; bear fewer than 10 non-informative (non-A/T/G/C, such as N or -) positions; and carry a non-empty CDR3 with no N and a nucleotide length that is a multiple of 3. For single-cell BCRs, cell-level quality control was also performed, requiring each cell to have either exactly one heavy chain and at least one light chain, or at least one heavy chain and exactly one light chain. Within a cell, for the chain type with more than one sequence, the most abundant sequence in terms of UMI count (when tied, the sequence that appeared earlier in the file) was kept. Ultimately, exactly one heavy chain and one light chain per cell were kept. Additionally, quality control against cross-sample contamination was performed by examining the extent, if any, of pairwise overlapping between samples in terms of BCRs with both identical UMIs and identical non-UMI nucleotide sequences.

Individualized genotypes were inferred based on sequences that passed all quality control using TIgGER v.1.0.0^41^ and used to finalize V(D)J annotations. Sequences annotated as non-productively rearranged by IgBLAST were removed from further analysis.

### Clonal lineage inference

B cell clonal lineages were inferred on a by-individual basis based on productively rearranged sequences using hierarchical clustering with single linkage^42^. When combining both bulk and single-cell BCRs, heavy chain-based clonal inference was performed^43^. First, heavy chain sequences were partitioned based on common V and J gene annotations and CDR3 lengths. Within each partition, heavy chain sequences with CDR3s that were within 0.15 normalized Hamming distance from each other were clustered as clones. When using only single-cell BCRs, clonal inference was performed based on paired heavy and light chains. First, paired heavy and light chains were partitioned based on common V and J gene annotations and CDR3 lengths. Within each partition, pairs whose heavy chain CDR3s were within 0.15 normalized Hamming distance from each other were clustered as clones.

Following clonal inference, full-length clonal consensus germline sequences were reconstructed for each clone with the D-segment (for heavy chains) and the N/P regions masked with Ns, resolving any ambiguous gene assignments by majority rule. Within each clone, duplicate IMGT-aligned V(D) J sequences from bulk sequencing were collapsed except for duplicates derived from different time points, tissues, B cell compartments, or isotypes.

### BCR analysis

BCR analysis was performed in R v4.1.0 with visualization performed using base R, ggplot2 v3.3.5^44^, and GraphPad Prism v9.

For the B cell compartment label, gene expression-based cluster annotation was used for single-cell BCRs; FACS-based sorting was used in general for bulk BCRs, except that PB sorts from lymph nodes were labelled LNPCs, d35 IgDlo sorts from blood were labelled activated, and d60 IgDlo sorts from blood were labelled memory. For the time point label, one blood PB sample that pooled collections on both d28 and d35 was treated as d28.

A heavy chain-based B cell clone was considered a S^+^ clone if the clone contained any sequence corresponding to a recombinant mAb that was synthesized based on the single-cell BCRs and that tested positive for S-binding.

Clonal overlap between B cell compartments was visualized using circlize v.0.4.13^45^.

Somatic hypermutation (SHM) frequency was calculated for each heavy chain sequence by counting the number of nucleotide mismatches from the germline sequence in the variable segment leading up to the CDR3, while excluding the first 18 positions that could be error-prone due to the primers used for generating the mAb sequences. Calculation was performed using the calcObservedMutations function from SHazaM v.1.0.2^40^.

Phylogenetic trees for S^+^ clones containing BMPCs were constructed on a by-participant basis using IgPhyML v1.1.3^46^ with the HLP19 model^47^. Only heavy chain sequences from the PB, GC, LNPC, RMB, and BMPC compartments were considered. For clones with >100 sequences, subsampling was applied with probabilities proportional to the proportions of sequences from different compartments, in addition to keeping all sequences corresponding to synthesized mAbs and all BMPC sequences. Only subsampled sequences from the PB, LNPC, and BMPC compartments were used for eventual tree-building. Trees were visualized using ggtree v3.0.4^48^.

### Human housekeeping genes

A list of human housekeeping genes was compiled from the 20 most stably expressed genes across 52 tissues and cell types in the Housekeeping and Reference Transcript (HRT) Atlas v1.0^49^; 11 highly uniform and strongly expressed genes reported^50^; and some of the most commonly used housekeeping genes^51^. The final list includes 34 genes: *ACTB, TLE5 (AES), AP2M1, BSG, C1orf43, CD59, CHMP2A, CSNK2B, EDF1, EEF2, EMC7, GABARAP, GAPDH, GPI, GUSB, HNRNPA2B1, HPRT1, HSP90AB1, MLF2, MRFAP1, PCBP1, PFDN5, PSAP, PSMB2, PSMB4, RAB11B, RAB1B, RAB7A, REEP5, RHOA, SNRPD3, UBC, VCP,* and *VPS29*.

### Processing of 10× Genomics single-cell 5′ gene expression data

Demultiplexed pair-end FASTQ reads were first preprocessed on a by-sample basis using the ‘cellranger count’ command from 10× Genomics’ Cell Ranger v.6.0.1 for alignment against the GRCh38 human reference v.2020-A (‘refdata-gex-GRCh38-2020-A’). To avoid a batch effect introduced by sequencing depth, the ‘cellranger aggr’ command was used to subsample from each sample so that all samples had the same effective sequencing depth, which was measured in terms of the number of reads confidently mapped to the transcriptome or assigned to the feature IDs per cell. Gene annotation on human reference chromosomes and scaffolds in Gene Transfer Format (‘gencode.v32.primary_assembly.annotation.gtf’) was downloaded (2021-06-02) from GENCODE v32^52^, from which a biotype (‘gene_type’) was extracted for each feature. Quality control was performed as follows on the aggregate gene expression matrix consisting of 360,803 cells and 36,601 features using SCANPY v1.7.2^53^ and Python v3.8.8. (1) To remove presumably lysed cells, cells with mitochondrial content greater than 12.5% of all transcripts were removed. (2) To remove likely doublets, cells with more than 8,000 features or 80,000 total UMIs were removed. (3) To remove cells with no detectable expression of common endogenous genes, cells with no transcript for any of the 34 housekeeping genes were removed. (4) The feature matrix was subset, based on their biotypes, to protein-coding, immunoglobulin, and T cell receptor genes that were expressed in at least 0.1% of the cells in any sample. The resultant feature matrix contained 15,744 genes. (5) Cells with detectable expression of fewer than 200 genes were removed. After quality control, there were a total of 318,156 cells from 47 single-cell samples (Extended Data Table 7).

### Single-cell gene expression analysis

Single-cell gene expression analysis was performed in SCANPY v1.7.2^53^. UMI counts measuring gene expression were log-normalized. The top 2,500 highly variable genes (HVGs) were identified using the ‘scanpy.pp.highly_variable_genes’ function with the ‘seurat_v3’ method, from which immunoglobulin and T cell receptor genes were removed. The data were scaled and centred, and principal component analysis (PCA) was performed based on HVG expression. PCA-guided neighborhood graphs embedded in Uniform Manifold Approximation and Projection (UMAP) were generated using the top 20 principal components via the ‘scanpy.pp.neighbors’ and ‘scanpy.tl.umap’ functions.

Overall clusters (Extended Data Table 5, top) were identified using Leiden graph-clustering via the ‘scanpy.tl.leiden’ function with resolution 0.15 (Extended Data Fig. 2b). UMAPs were faceted by batch, by participant, and by participant followed by sample; and inspected for convergence across batches, participants, and samples within participants, to assess whether there was a need for integration (Extended Data Fig. 2b). Cluster identities were assigned by examining the expression of a set of marker genes for different cell types (Extended Data Fig. 2c): *MS4A1*, *CD19* and *CD79A* for B cells; *CD3D, CD3E, CD3G, IL7R* and *CD4* or *CD8A* for CD4^+^ or CD8^+^ T cells, respectively; *GZMB, GNLY, NKG7* and *NCAM1* for natural killer (NK) cells; *CD14, LYZ, CST3* and *MS4A7* for monocytes; *IL3RA* and *CLEC4C* for plasmacytoid dendritic cells (pDCs); and *FDCSP, CXCL14*^20^ and *FCAMR*^21^ for follicular dendritic cells (FDCs). To remove potential contamination by platelets, 60 cells with a log-normalized expression value of >2.5 for PPBP were removed. All 349 cells from the FDC cluster were confirmed to have originated from FNA samples instead of blood.

Cells from the overall B cell cluster (Extended Data Table 5, bottom) were further clustered to identify B cell subsets using Leiden graph-clustering via the ‘scanpy.tl.leiden’ function with resolution 0.2 (Extended Data Fig. 2d). Cluster identities were assigned by examining the expression of a set of marker genes for different B cell subsets (Extended Data Fig. 2e) along with the availability of BCRs. The following marker genes were examined: *BCL6, RGS13, MEF2B, STMN1, ELL3* and *SERPINA9* for GC B cells; *XBP1, IRF4, SEC11C, FKBP11, JCHAIN* and *PRDM1* for PBs and LNPCs; *TCL1A, IL4R, CCR7, IGHM,* and *IGHD* for naive B cells; and *TNFRSF13B, CD27* and *CD24* for RMB cells. Although 2 groups clustered with B cells during overall clustering, they were labelled ‘B and T’ as their cells tended to have both BCRs and high expression levels of *CD2* and *CD3E*; and were subsequently excluded from the final B cell clustering. 19 cells that were found in the GC cluster but came from blood samples were labelled ‘PB-like’. 198 cells that were found in the PB cluster but came from FNA samples were re-assigned as LNPCs. 37 cells that were found in the LNPC cluster but came from blood samples were re-assigned as PBs. Heavy chain SHM frequency and isotype usage of the B cell subsets were assessed for consistency with expected values to further confirm their assigned identities.

### Selection and curation of single-cell BCRs from GC/LNPC for expression

Single-cell gene expression analysis was performed using lymph node samples on a by-participant basis. Clonal inference was performed based on paired heavy and light chains from the same samples. From every clone with a clone size of >3 cells that contained cells from the GC and/or LNPC clusters, one GC or LNPC cell was selected. For selection, where a clone spanned both the GC and LNPC compartments, and/or multiple time points, a compartment and a timepoint were first randomly selected. Within that clone, the cell with the highest heavy chain UMI count was then selected, breaking ties based on *IGHV* SHM frequency. In all selected cells, native pairing was preserved.

BCRs from the selected cells were curated prior to synthesis. First, artificial gaps introduced under the IMGT unique numbering system^54^ were removed from the IMGT-aligned observed V(D)J sequences. IMGT gaps were identified as positions containing in-frame triplet dots (‘…’) in the IMGT-aligned germline sequences. Second, any non-informative (non-A/T/G/C, such as N or −) positions in the observed sequences, with the exception of potential in-frame indels, were patched by the nucleotides at their corresponding germline positions. Third, if applicable, the 3’ end of the observed sequences were trimmed so that the total nucleotide length would be a multiple of 3. Finally, potential in-frame indels were manually reviewed. For a given potential in-frame indel from a selected cell, its presence or lack thereof in the unselected cells from the same clone was considered. Barring strong indications that an in-frame indel was due to sequencing error rather than the incapability of the IMGT unique numbering system of capturing it, the in-frame indels were generally included in the final curated sequences.

### Transfection for recombinant mAbs production

Selected pairs of heavy and light chain sequences were synthesized by GenScript and sequentially cloned into IgG1 and Igκ/λ expression vectors, respectively. Heavy and light chain plasmids were co-transfected into Expi293F cells (Thermo Fisher Scientific) for recombinant mAb production, followed by purification with protein A agarose resin (GoldBio). Expi293F cells were cultured in Expi293 Expression Medium (Gibco) according to the manufacturer’s protocol.

## Acknowledgements

We thank the generous participation of the donors for providing specimens. The Ellebedy laboratory was supported by National Institute of Allergy and Infectious Diseases (NIAID) grants U01AI141990 and 1U01AI150747, NIAID Centers of Excellence for Influenza Research and Surveillance contract HHSN272201400006C and HHSN272201400008C, and NIAID Collaborative Influenza Vaccine Innovation Centers contract 75N93019C00051. W.K. was supported by the Basic Science Research Program through the National Research Foundation of Korea (NRF) funded by the Ministry of Education (2021R1A6A3A03041509). J.S.T. was supported by NIAID 5T32CA009547. This study utilized samples obtained from the Washington University School of Medicine’s COVID-19 biorepository supported by the NIH/National Center for Advancing Translational Sciences, grant number UL1 TR002345. The WU353 and WU368 studies were reviewed and approved by the Washington University Institutional Review Board (approval no. 202003186 and 202012081, respectively).

## Author Contributions

A.H.E. conceived and designed the study. A.H., M.K.K., L.P., P.A.M., I.P., J.A.O., and R.M.P. wrote and maintained the IRB protocol, recruited, and phlebotomized participants, coordinated sample collection, and analyzed clinical data. W.K., E.K, and J.S.T. processed specimens. W.K., E.K., W.B.A., and J.S.T. performed ELISA and ELISpot. W.K., S.C.H, A.J.Schmitz, T.L., M.T., and W.B.A. generated and characterized monoclonal antibodies. W.K.and A.J.Sturtz prepared libraries for scRNA sequencing. A.J.Schmitz performed RNA extractions and library preparation for BCR bulk sequencing. J.Q.Z. analyzed scRNA sequencing and BCR bulk sequencing data. A.J.Schmitz expressed SARS-CoV-2 S and RBD proteins. J.S.T. sorted cells and collected and analysed the flow cytometry data. T.S. and W.D.M performed FNA. W.D.M. and S.A.T. supervised lymph node evaluation prior to FNA and specimen collection and evaluated lymph node ultrasound data. W.K., J.Q.Z, J.S.T., and A.H.E. analyzed the data. A.H.E. supervised experiments and obtained funding. W.K., P.A.M, J.S.T., and A.H.E. composed the manuscript. All authors reviewed the manuscript.

## Competing Interests

The Ellebedy laboratory received funding under sponsored research agreements that are unrelated to the data presented in the current study from Emergent BioSolutions and from AbbVie. A.H.E. is a consultant for Mubadala Investment Company and the founder of ImmuneBio Consulting. J.S.T. is a consultant for Gerson Lehrman Group. J.S.T., A.J.Schmitz., and A.H.E. are recipients of a licensing agreement with Abbvie that is unrelated to the data presented in the current study. A patent application related to this work has been filed by Washington University School of Medicine. The content of this manuscript is solely the responsibility of the authors and does not necessarily represent the official view of NIAID or NIH.

## Data availability statement

New raw sequencing data, processed transcriptomics data, and processed BCR data will be uploaded to Sequence Read Archive, GEO, and Zenodo respectively before final publication. Previously reported bulk-sequenced BCR data used in this study were deposited under PRJNA731610 and PRJNA741267 on SRA, and at https://doi.org/10.5281/zenodo.5042252 and https://doi.org/10.5281/zenodo.5040099 on Zenodo.

## Extended Data Figure legends

**Extended Data Figure 1. Persistence of humoral immune responses to SARS-CoV-2 mRNA vaccination. a**, Flow cytometry gating strategies for GC B cells (Fig. 1b) and LNPCs (defined as CD19^+^ CD3^−^ IgD^low^ CD20^low^ CD38^+^ BLIMP1^+^ CD71^+^ live singlet lymphocytes) in the lymph node. **b**, Kinetics of total (left) and S^+^ LNPCs (right) as gated in **a**. **c**, Frequencies of BMPCs secreting IgA antibodies specific for the indicated antigens 29 weeks after immunization. **d**, **e**, Plasma antibody titers against SARS-CoV-2 S measured by ELISA in participants without (red) and with (black) a history of SARS-CoV-2 infection in SARS-CoV-2 vaccinated (left, center) and unvaccinated (right) participants 29 weeks after the first vaccine dose or symptom onset (**d**) and in vaccinated participants over time (**e**). *P* values were determined by Kruskal-Wallis test followed by Dunn’s multiple comparison test between unvaccinated and both vaccinated groups (**d**), and by two-sided Mann-Whitney test (**e**). Horizontal lines indicate median values in **c** and **e**. **f**, Flow cytometry gating strategies for MBCs (CD19^+^ CD3^−^ IgD^low^ CD20^+^ CD38^−^ live singlet lymphocytes) and S^+^ MBCs (Fig. 1g) in blood.

**Extended Data Figure 2. Identification of SARS-CoV-2 S-binding B cell clones in the lymph node. a**, Flow cytometry gating strategies for sorting PBs (defined as CD19^+^ CD3^−^ IgD^low^ CD20^low^ CD38^+^ CD71^+^ live singlet lymphocytes) from blood. **b**, **d** UMAPs of scRNA-seq data from PBs sorted from blood and FNA of draining axillary lymph nodes (b), and UMAPs of B cell scRNA-seq clusters (d). **c, e,** Dot plots for the marker genes used for identifying annotated clusters. **f**, Flow cytometry gating strategies for sorting GC B cells (CD19^+^ CD4^−^ IgD^low^ CD20^+^ CD38^int^ CXCR5^high^ CD71^+^ live singlet lymphocytes) and LNPCs (CD19^+^ CD4^−^ IgD^low^ CD20^low^ CD38^+^ CXCR5^low^ CD71^+^ live singlet lymphocytes) from FNAs. **g**, Visualization of SARS-CoV-2 S-binding clones from all participants on integrated UMAP of B cell clusters. Percentages are of SARS-CoV-2 S-binding clones within GC B cells (blue), LNPCs (green), PBs (red), RMBs (pink) or naive B cells (yellow).

**Extended Data Figure 3. Maturation of SARS-CoV-2 S-binding B cells in the lymph node. a**, Comparison of *IGHV* nucleotide mutation frequency of SARS-CoV-2 S-binding GC B cells in each participant at the indicated time points. Horizontal lines represent median values.

**Extended Data Figure 4. Evolution of B cell clones induced by SARS-CoV-2 vaccination. a**, Comparison of *IGHV* nucleotide mutation frequency of each B cell subsets. Horizontal lines represent median values. *P* values are determined by Kruskal-Wallis test followed by Dunn’s multiple comparison test.

**Extended Data Table legends**

**Extended Data Table 1. Demographics of participants**

**Extended Data Table 2. Vaccine side effects**

**Extended Data Table 3. Frequencies of total and SARS-CoV-2 S-binding GC B cells and LNPCs in draining axillary lymph nodes**

**Extended Data Table 4. Frequencies of CD14^+^ cells in lymph node samples**

**Extended Data Table 5. Cell counts and frequencies of overall and B cell clusters in scRNA-seq of PBs sorted from blood and FNA from lymph nodes**

**Extended Data Table 6. Description of recombinant mAbs derived from GC B cells and LNPCs**

**Extended Data Table 7. Processing of BCR and 5’ gene expression data from scRNA-seq**

**Extended Data Table 8. Processing of BCR reads from bulk-seq**

## Notes

https://doi.org/10.5281/zenodo.5042252

https://doi.org/10.5281/zenodo.5040099

