## Supplemental Figures and Tables for "Germinal centre-driven maturation of B cell response to SARS-CoV-2 vaccination"

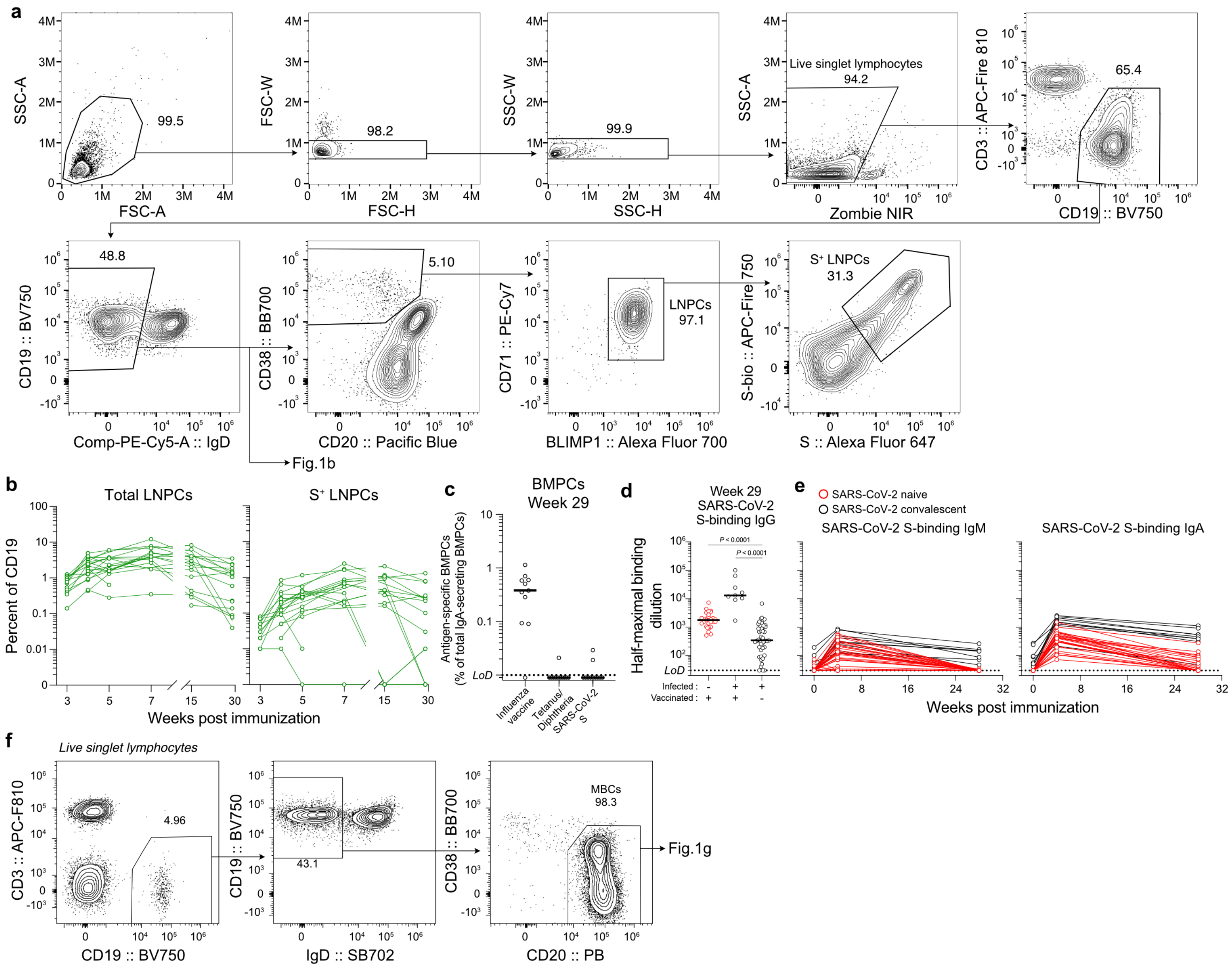


**Extended Data Figure 1. Persistence of humoral immune responses to SARS-CoV-2 mRNA vaccination. a**, Flow cytometry gating strategies for GC B cells (Fig. 1b) and LNPCs (defined as $CD19^+ CD3^- IgD^{low} CD20^{low} CD38^+ BLIMP1^+ CD71^+$ live singlet lymphocytes) in the lymph node. **b**, Kinetics of total (left) and $S^+$ LNPCs (right) as gated in **a**. **c**, Frequencies of BMPCs secreting IgA antibodies specific for the indicated antigens 29 weeks after immunization. **d**, **e**, Plasma antibody titers against SARS-CoV-2 S measured by ELISA in participants without (red) and with (black) a history of SARS-CoV-2 infection in SARS-CoV-2 vaccinated (left, center) and unvaccinated (right) participants 29 weeks after the first vaccine dose or symptom onset (**d**) and in vaccinated participants over time (**e**) . *P* values were determined by Kruskal-Wallis test followed by Dunn's multiple comparison test between unvaccinated and both vaccinated groups (**d**), and by two-sided Mann-Whitney test (**e**). Horizontal lines indicate median values in **c** and **e**. **f**, Flow cytometry gating strategies for MBCs ($CD19^+ CD3^- IgD^{low} CD20^+ CD38^-$ live singlet lymphocytes) and $S^+$ MBCs (Fig. 1g) in blood.

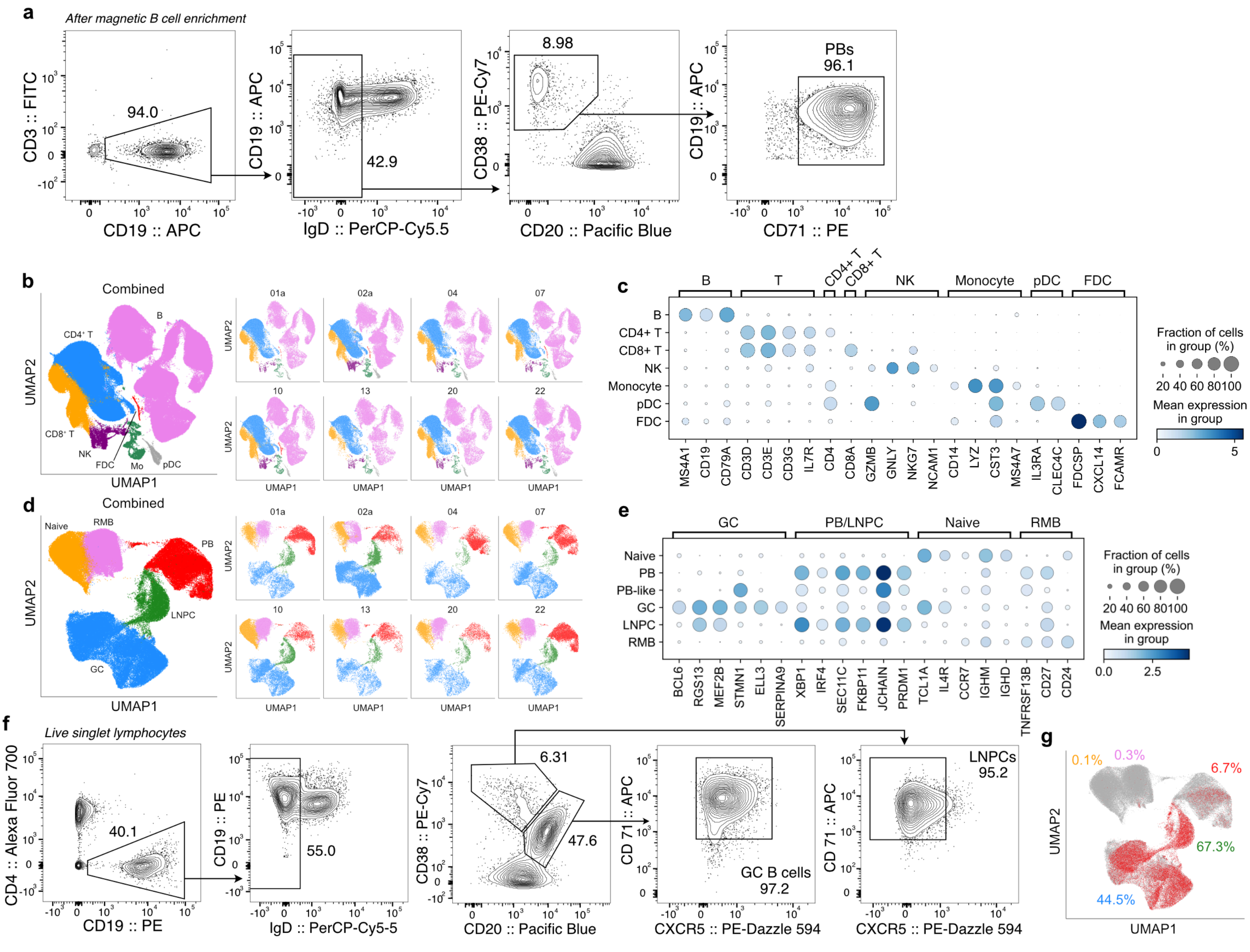


**Extended Data Figure 2. Identification of SARS-CoV-2 S-binding B cell clones in the lymph node. a**, Flow cytometry gating strategies for sorting PBs (defined as CD19$^{+}$ CD3$^{-}$ IgD$^{low}$ CD20$^{low}$ CD38$^{+}$ CD71$^{+}$ live singlet lymphocytes) from blood. **b**, **d** UMAPs of scRNA-seq data from PBs sorted from blood and FNA of draining axillary lymph nodes (b), and UMAPs of B cell scRNA-seq clusters (d). **c, e,** Dot plots for the marker genes used for identifying annotated clusters. **f**, Flow cytometry gating strategies for sorting GC B cells (CD19$^{+}$ CD4$^{-}$ IgD$^{low}$ CD20$^{+}$ CD38$^{int}$ CXCR5$^{high}$ CD71$^{+}$ live singlet lymphocytes) and LNPCs (CD19$^{+}$ CD4$^{-}$ IgD$^{low}$ CD20$^{low}$ CD38$^{+}$ CXCR5$^{low}$ CD71$^{+}$ live singlet lymphocytes) from FNAs. **g**, Visualization of SARS-CoV-2 S-binding clones from all participants on integrated UMAP of B cell clusters. Percentages are of SARS-CoV-2 S-binding clones within GC B cells (blue), LNPCs (green), PBs (red), RMBs (pink) or naive B cells (yellow).

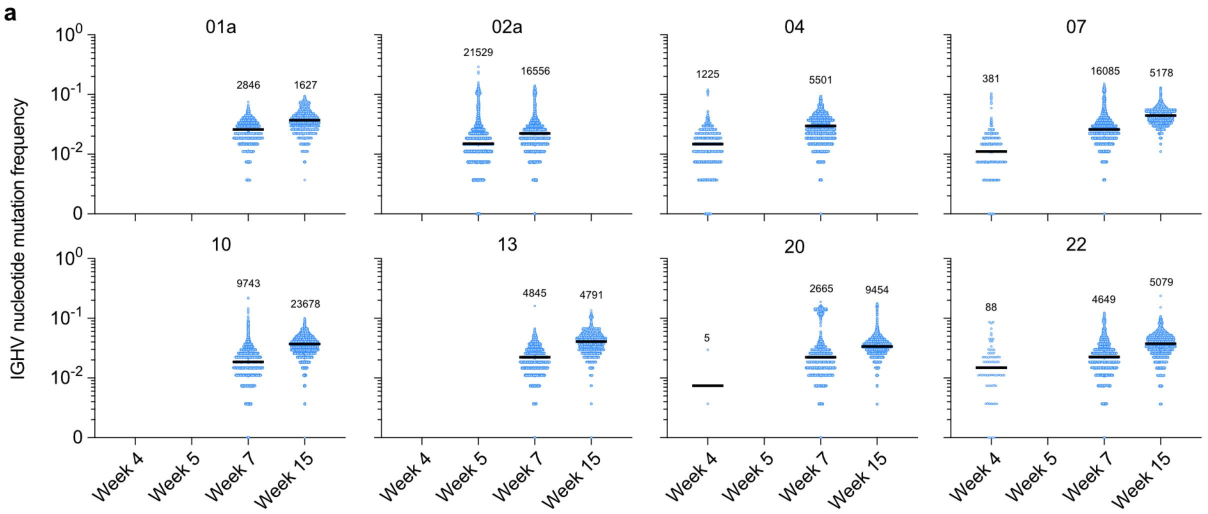


**Extended Data Figure 3. Maturation of SARS-CoV-2 S-binding B cells in the lymph node.** **a**, Comparison of *IGHV* nucleotide mutation frequency of SARS-CoV-2 S-binding GC B cells in each participant at the indicated time points. Horizontal lines represent median values.

a

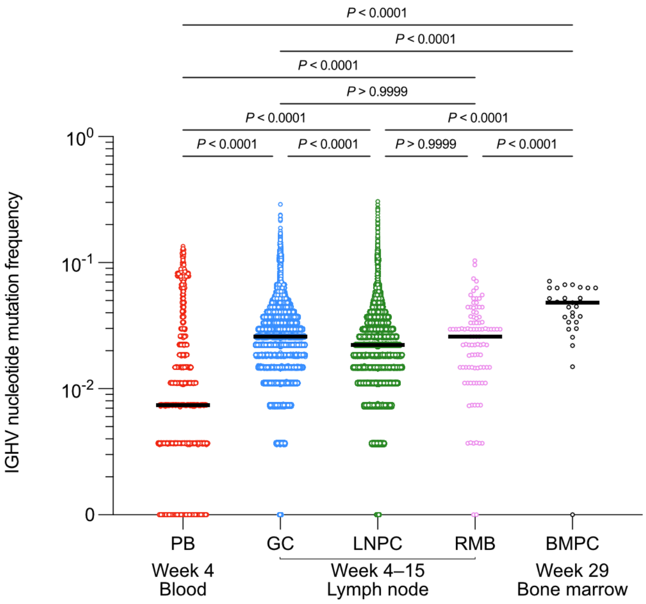


**Extended Data Figure 4. Evolution of B cell clones induced by SARS-CoV-2 vaccination.** **a**, Comparison of *IGHV* nucleotide mutation frequency of each B cell subsets. Horizontal lines represent median values. *P* values are determined by Kruskal-Wallis test followed by Dunn's multiple comparison test.

### Extended Data Table 1. Demographics of participants

| Variable | SARS-CoV-2 mRNA vaccination Total (N=43) N (%) | Blood (N=42) N (%) | Lymph node (N=15) N (%) | Bone marrow (N=11) N (%) | Variable | Convalescent (N=48) N (%) |
|---|---|---|---|---|---|---|
| **Age (median [range])** | 38 (28-73) | 37.5 (28-73) | 37 (28-52) | 36 (28-48) | **Age (median [range])** | 50 (21-69) |
| **Sex** | | | | | **Sex** | |
| Female | 21 (48.8) | 20 (47.6) | 7 (53.8) | 6 (54.5) | Female | 23 (47.9) |
| Male | 22 (51.2) | 22 (52.4) | 6 (46.2) | 5 (45.5) | Male | 25 (52.1) |
| **Race** | | | | | **Race/Ethnicity** | |
| White | 34 (79.1) | 34 (81) | 12 (80) | 11 (100) | White | 47 (97.9) |
| Black | 6 (14) | 6 (14.3) | 2 (13.3) | 0 (0) | Black | 0 (0) |
| Asian | 1(2.3) | 0 (0) | 1 (6.7) | 0 (0) | Asian | 1 (2.1) |
| Other | 2 (4.7) | 2 (4.8) | 0 (0) | 0 (0) | Hispanic | 0 (0) |
| **Ethnicity** | | | | | | |
| Not of Hispanic, Latinx, or Spanish origin | 41 (95.3) | 40 (95.2) | 14 (93.3) | 10 (90.9) | | |
| Hispanic, Latinx, Spanish origin | 2 (4.7) | 2 (4.8) | 1 (6.7) | 1 (9.1) | | |
| **BMI (media [range])** | 26.8 (21.4-67.4) | 26.9 (21.4-67.4) | 24.1 (21.4-40.1) | 23.9 (21.4-40.1) | | |
| **Comorbidities** | | | | | **Comorbidities** | |
| Lung disease | 1 (2.3) | 1 (2.4) | 0 (0) | 0 (0) | Asthma | 10 (20.8) |
| Diabetes mellitus | 0 (0) | 0 (0) | 0 (0) | 0 (0) | Other lung disease | 0 (0) |
| Hypertension | 7 (16.3) | 6 (14.3) | 2 (13.3) | 1 (9.1) | Heart disease | 0 (0) |
| Cardiovascular | 0 (0) | 0 (0) | 0 (0) | 0 (0) | Hypertension | 7 (14.6) |
| Liver disease | 0 (0) | 0 (0) | 0 (0) | 0 (0) | Diabetes mellitus | 3 (6.3) |
| Chronic kidney disease | 0 (0) | 0 (0) | 0 (0) | 0 (0) | Cancer | 6 (12.5) |
| Cancer on chemotherapy | 0 (0) | 0 (0) | 0 (0) | 0 (0) | Autoimmune disease | 4 (8.3) |
| Hematological malignancy | 0 (0) | 0 (0) | 0 (0) | 0 (0) | Hyperlipidaemia | 2 (4.2) |
| Pregnancy | 0 (0) | 0 (0) | 0 (0) | 0 (0) | GERD | 5 (10.4) |
| Neurological | 0 (0) | 0 (0) | 0 (0) | 0 (0) | Other | 16 (33.3) |
| HIV | 0 (0) | 0 (0) | 0 (0) | 0 (0) | Solid Organ Transplant | 1 (2.1) |
| Hyperlipidemia | 1 (2.3) | 1 (2.4) | 0 (0) | 0 (0) | | |
| Gastroesophageal reflux disease | - | - | - | - | | |
| Autoimmune disease | - | - | - | - | | |
| Solid organ transplant recipient | 0 (0) | 0 (0) | 0 (0) | 0 (0) | | |
| Bone marrow transplant recipient | 0 (0) | 0 (0) | 0 (0) | 0 (0) | | |
| **Confirmed SARS-CoV-2 infection** | 13 (30.2) | 13 (31) | 0 (0) | 0 (0) | **Hospitalized for treatment of COVID-19** | 3 (6.3) |
| Time from SARS-CoV-2 infection to baseline visit in days (median [range]) | 234 (70-370) | 234 (70-370) | - | - | | |

#### Extended Data Table 2. Vaccine side effects

| Variable | Total N=43<br>N (%) | Total N=43<br>N (%) |
|---|---|---|
| | **First dose** | **Second dose** |
| None | 2 (4.7) | 0 (0) |
| Chills | 9 (20.9) | 18 (41.9) |
| Fever | 6 (14) | 12 (27.9) |
| Headache | 10 (23.3) | 16 (37.2) |
| Injection site pain/redness/swelling | 37 (86) | 39 (90.7) |
| Muscle or joint pain | 10 (23.3) | 25 (58.1) |
| Fatigue | 13 (30.2) | 27 (62.8) |
| | **Duration of side effects in hours (median [range])** | |
| Chills | 18 (6-72) | 24 (0.2-48) |
| Fever | 8 (3-72) | 30 (1-48) |
| Headache | 36 (6-120) | 24 (4-72) |
| Injection site pain | 48 (0.2-168) | 48 (2-144) |
| Muscle or joint pain | 24 (3-72) | 24 (1-48) |
| Fatigue | 48 (12-120) | 24 (3-144) |

#### Extended Data Table 3. Frequencies of total and SARS-CoV-2 S-binding GC B cells and LNPCs in draining axillary lymph nodes

| Participant | LN # | Total GC B cells (% of CD19) | | | | | | SARS-CoV-2 S-binding GC B cells (% of CD19) | | | | | |
|---|---|---|---|---|---|---|---|---|---|---|---|---|---|
| | | Week 3 | Week 4 | Week 5 | Week 7 | Week 15 | Week 29 | Week 3 | Week 4 | Week 5 | Week 7 | Week 15 | Week 29 |
| 01a | 1 | 15.25 | 13.69 | 11.14 | 31.67 | 29.13 | 14.37 | 3.23 | 5.41 | 4.01 | 13.75 | 18.06 | 8.00 |
| 02a | 1 | 8.28 | 34.16 | 44.92 | 21.90 | 7.17 | 0.85 | 1.36 | 9.66 | 11.55 | 5.98 | 0.01 | 0.06 |
| | 2 | | 14.08 | 13.54 | 23.13 | | | | 4.66 | 3.85 | 9.33 | | |
| 04 | 1 | 4.71 | 14.02 | 11.21 | 43.94 | 3.75 | 0.38 | 0.57 | 2.84 | 5.59 | 6.37 | 0.70 | 0.00 |
| | 2 | | | | | 1.09 | 0.41 | | | | | 0.03 | 0.00 |
| 07 | 1 | 21.14 | 19.68 | 11.48 | 39.22 | 28.93 | 28.10 | 4.59 | 4.92 | 3.87 | 14.55 | 19.92 | 14.77 |
| 08 | 1 | 9.91 | 1.31 | 3.99 | 12.15 | | 31.34 | 1.21 | 0.41 | 0.91 | 4.21 | | 23.33 |
| 10 | 1 | 7.72 | 5.92 | 2.82 | 7.34 | 19.33 | 19.41 | 1.10 | 1.47 | 0.90 | 3.10 | 14.19 | 15.37 |
| | 2 | | 5.70 | 3.40 | 4.28 | 7.14 | 9.19 | | 1.36 | 1.04 | 1.05 | 2.79 | 5.90 |
| 13 | 1 | 14.99 | 8.69 | 7.23 | 16.02 | 20.21 | 0.77 | 3.53 | 3.00 | 2.14 | 3.70 | 10.03 | 0.00 |
| 15 | 1 | 13.06 | 26.89 | 44.56 | | | | 2.96 | 8.01 | 9.58 | | | |
| | 2 | | 0.23 | 0.61 | | | | | 0.01 | 0.02 | | | |
| 16 | 1 | 5.20 | 7.61 | 17.04 | | 5.82 | 4.40 | 0.74 | 1.50 | 3.34 | | 2.57 | 1.48 |
| | 2 | | 1.22 | 7.44 | | 9.61 | 3.64 | | 0.24 | 1.84 | | 5.81 | 1.06 |
| 20 | 1 | 0.41 | 7.49 | 2.10 | 0.52 | 0.47 | 0.71 | 0.03 | 3.07 | 0.31 | 0.04 | 0.04 | 0.02 |
| | 2 | | 7.79 | 3.36 | 13.67 | 16.93 | 20.23 | | 1.28 | 0.72 | 4.43 | 11.08 | 10.56 |
| 21 | 1 | | 20.18 | | 14.50 | | 7.03 | | 5.92 | | 5.20 | | 3.33 |
| 22 | 1 | 24.69 | 20.63 | 19.66 | 25.86 | 22.01 | 0.72 | 4.44 | 5.35 | 4.92 | 7.33 | 9.69 | 0.00 |
| 26 | 1 | | | | | | 1.69 | | | | | | 0.29 |
| 28 | 1 | 6.40 | 6.25 | | | 12.82 | 9.46 | 1.12 | 1.16 | | | 7.26 | 4.73 |
| 43 | 1 | 29.01 | 29.38 | 26.09 | 29.43 | 35.19 | | 4.43 | 9.42 | 6.28 | 15.26 | 15.38 | |

| Participant | LN # | Total LNPCs (% of CD19) | | | | | | SARS-CoV-2 S-binding LNPCs (% of CD19) | | | | | |
|---|---|---|---|---|---|---|---|---|---|---|---|---|---|
| | | Week 3 | Week 4 | Week 5 | Week 7 | Week 15 | Week 29 | Week 3 | Week 4 | Week 5 | Week 7 | Week 15 | Week 29 |
| 01a | 1 | 0.77 | 4.96 | 6.00 | 6.82 | 8.23 | 2.33 | 0.07 | 0.84 | 1.15 | 1.61 | 2.04 | 0.78 |
| 02a | 1 | 0.35 | 2.47 | 3.52 | 3.20 | 0.43 | 0.22 | 0.01 | 0.20 | 0.40 | 0.36 | 0.00 | 0.00 |
| | 2 | | 2.30 | 3.74 | 3.91 | | | | 0.25 | 0.28 | 0.54 | | |
| 04 | 1 | 0.88 | 1.29 | 1.57 | 2.67 | 0.29 | 0.08 | 0.05 | 0.09 | 0.43 | 0.20 | 0.00 | 0.01 |
| | 2 | | | | | 0.17 | 0.14 | | | | | 0.00 | 0.01 |
| 07 | 1 | 0.97 | 2.74 | 1.73 | 3.97 | 2.02 | 1.87 | 0.07 | 0.26 | 0.26 | 0.67 | 0.32 | 0.22 |
| 08 | 1 | 1.07 | 2.18 | 1.53 | 4.84 | | 3.49 | 0.07 | 0.09 | 0.15 | 0.74 | | 1.30 |
| 10 | 1 | 0.71 | 4.08 | 2.31 | 3.92 | 4.02 | 2.42 | 0.04 | 0.34 | 0.33 | 0.86 | 1.06 | 0.76 |
| | 2 | | 2.83 | 3.51 | 1.75 | 1.08 | 1.40 | | 0.18 | 0.40 | 0.16 | 0.19 | 0.31 |
| 13 | 1 | 0.39 | 0.94 | 1.57 | 5.25 | 3.02 | 0.04 | 0.02 | 0.21 | 0.30 | 0.64 | 0.44 | 0.00 |
| 15 | 1 | 1.26 | 1.27 | 3.60 | | | | 0.08 | 0.12 | 0.29 | | | |
| | 2 | | 1.79 | 0.64 | | | | | 0.10 | 0.01 | | | |
| 16 | 1 | 0.47 | 0.88 | 2.24 | | 1.68 | 0.60 | 0.02 | 0.05 | 0.12 | | 0.16 | 0.03 |
| | 2 | | 0.43 | 1.31 | | 2.91 | 1.06 | | 0.02 | 0.11 | | 0.43 | 0.10 |
| 20 | 1 | 0.14 | 0.46 | 0.28 | 0.35 | 0.34 | 0.09 | 0.01 | 0.01 | 0.00 | 0.00 | 0.00 | 0.00 |
| | 2 | | 1.73 | 1.62 | 4.01 | 2.48 | 1.32 | | 0.05 | 0.08 | 0.22 | 0.42 | 0.29 |
| 21 | 1 | | 3.42 | | 12.18 | | 1.53 | | 0.38 | | 1.70 | | 0.25 |
| 22 | 1 | 1.01 | 2.92 | 5.91 | 7.84 | 5.84 | 0.07 | 0.03 | 0.16 | 0.30 | 0.77 | 0.82 | 0.00 |
| 26 | 1 | | | | | | 0.23 | | | | | | 0.00 |
| 28 | 1 | 0.54 | 1.54 | | | 6.26 | 1.29 | 0.03 | 0.13 | | | 1.15 | 0.25 |
| 43 | 1 | 1.14 | 3.85 | 3.66 | 7.75 | 3.63 | | 0.05 | 0.63 | 0.55 | 2.44 | 0.93 | |

#### Extended Data Table 4. Frequencies of $CD14^{+}$ cells in lymph node samples

| Participant | LN # | CD14 (% of live singlet) | | | | | |
|---|---|---|---|---|---|---|---|
| | | Week 3 | Week 4 | Week 5 | Week 7 | Week 15 | Week 29 |
| 01a | 1 | 0.21 | 0.06 | 0.15 | 0.15 | 0.16 | 0.21 |
| 02a | 1 | 0.16 | 0.17 | 0.07 | 0.25 | 0.17 | 0.30 |
| | 2 | | 0.14 | 0.14 | 0.71 | | |
| 04 | 1 | 0.19 | 0.37 | 0.72 | 0.14 | 1.00 | 0.16 |
| | 2 | | | | | 0.33 | 0.25 |
| 07 | 1 | 0.19 | 0.15 | 0.42 | 0.66 | 0.04 | 4.38 |
| 08 | 1 | 0.27 | 0.17 | 0.34 | 0.81 | | 1.01 |
| 10 | 1 | 0.22 | 0.07 | 0.08 | 0.21 | 0.03 | 0.23 |
| | 2 | | 0.09 | 0.05 | 0.13 | 0.03 | 0.26 |
| 13 | 1 | 0.57 | 0.34 | 0.43 | 0.33 | 1.63 | 1.04 |
| 15 | 1 | 0.07 | 0.19 | 0.08 | | | |
| | 2 | | 0.14 | 0.24 | | | |
| 16 | 1 | 0.26 | 0.14 | 0.06 | | 1.21 | 0.50 |
| | 2 | | 0.11 | 0.14 | | 0.51 | 0.32 |
| 20 | 1 | 0.21 | 0.15 | 0.29 | 0.41 | 0.84 | 0.11 |
| | 2 | | 0.11 | 0.16 | 0.11 | 0.29 | 0.16 |
| 21 | 1 | | 0.13 | | 0.11 | | 0.51 |
| 22 | 1 | 0.18 | 0.16 | 0.20 | 0.36 | 0.28 | 0.24 |
| 26 | 1 | | | | | | 0.03 |
| 28 | 1 | 0.14 | 0.22 | | | 0.66 | 0.66 |
| 43 | 1 | 0.52 | 0.32 | 0.98 | 0.09 | 0.01 | |

#### Extended Data Table 5. Cell counts and frequencies of overall and B cell clusters in scRNA-seq of PBs sorted from blood and FNA from lymph nodes

| Sample | Overall cluster | Cell count (% of whole cells) |
|---|---|---|
| PBs sorted from blood | B | 27420 (96.5%) |
| | CD4+ T | 35 (0.1%) |
| | CD8+ T | 87 (0.3%) |
| | NK | 730 (2.6%) |
| | Monocyte | 142 (0.5%) |
| | pDC | 6 (0%) |
| | FDC | 0 (0%) |
| Lymph node | B | 139287 (48.1%) |
| | CD4+ T | 108647 (37.5%) |
| | CD8+ T | 29986 (10.4%) |
| | NK | 4983 (1.7%) |
| | Monocyte | 4260 (1.5%) |
| | pDC | 2164 (0.7%) |
| | FDC | 349 (0.1%) |
| Combined | B | 166707 (52.4%) |
| | CD4+ T | 108682 (34.2%) |
| | CD8+ T | 30073 (9.5%) |
| | NK | 5713 (1.8%) |
| | Monocyte | 4402 (1.4%) |
| | pDC | 2170 (0.7%) |
| | FDC | 349 (0.1%) |

| Participants | B cell cluster | Total cells (each B cell cluster out of whole B cells) | SARS-CoV-2 S-binding cells (S-binding cells out of total cells within each cluster) |
|---|---|---|---|
| 01a | GC | 4050 (25.5%) | 2258 (55.8%) |
| | LNPC | 1113 (7%) | 891 (80.1%) |
| | Naïve | 2034 (12.8%) | 0 (0%) |
| | PB | 6183 (38.9%) | 424 (6.9%) |
| | PB-like | 2 (0%) | 0 (0%) |
| | RMB | 2495 (15.7%) | 15 (0.6%) |
| 02a | GC | 17869 (56.5%) | 7663 (42.9%) |
| | LNPC | 3101 (9.8%) | 2285 (73.7%) |
| | Naïve | 4161 (13.2%) | 2 (0%) |
| | PB | 1837 (5.8%) | 246 (13.4%) |
| | PB-like | 1 (0%) | 0 (0%) |
| | RMB | 4671 (14.8%) | 32 (0.7%) |
| 04 | GC | 3694 (30%) | 1602 (43.4%) |
| | LNPC | 392 (3.2%) | 109 (27.8%) |
| | Naïve | 2059 (16.7%) | 1 (0%) |
| | PB | 4363 (35.5%) | 212 (4.9%) |
| | PB-like | 0 (0%) | - |
| | RMB | 1788 (14.5%) | 5 (0.3%) |
| 07 | GC | 6313 (37.1%) | 3695 (58.5%) |
| | LNPC | 716 (4.2%) | 492 (68.7%) |
| | Naïve | 3314 (19.5%) | 0 (0%) |
| | PB | 3667 (21.6%) | 160 (4.4%) |
| | PB-like | 9 (0.1%) | 0 (0%) |
| | RMB | 2989 (17.6%) | 10 (0.3%) |
| 10 | GC | 8505 (24.2%) | 3099 (36.4%) |
| | LNPC | 2419 (6.9%) | 1424 (58.9%) |
| | Naïve | 13382 (38%) | 13 (0.1%) |
| | PB | 1497 (4.3%) | 59 (3.9%) |
| | PB-like | 2 (0%) | 0 (0%) |
| | RMB | 9365 (26.6%) | 18 (0.2%) |
| 13 | GC | 3635 (29.7%) | 1385 (38.1%) |
| | LNPC | 907 (7.4%) | 671 (74%) |
| | Naïve | 2637 (21.5%) | 6 (0.2%) |
| | PB | 1861 (15.2%) | 38 (2%) |
| | PB-like | 3 (0%) | 0 (0%) |
| | RMB | 3211 (26.2%) | 1 (0%) |
| 20 | GC | 3256 (27.2%) | 1641 (50.4%) |
| | LNPC | 659 (5.5%) | 363 (55.1%) |
| | Naïve | 3100 (25.9%) | 0 (0%) |
| | PB | 1178 (9.9%) | 23 (2%) |
| | PB-like | 0 (0%) | - |
| | RMB | 3766 (31.5%) | 3 (0.1%) |
| 22 | GC | 5281 (26.9%) | 2087 (39.5%) |
| | LNPC | 1576 (8%) | 1089 (69.1%) |
| | Naïve | 2072 (10.6%) | 0 (0%) |
| | PB | 6633 (33.8%) | 660 (10%) |
| | PB-like | 2 (0%) | 0 (0%) |
| | RMB | 4036 (20.6%) | 14 (0.3%) |
| Combined | GC | 52603 (33.8%) | 23430 (44.5%) |
| | LNPC | 10883 (7%) | 7324 (67.3%) |
| | Naïve | 32759 (21%) | 22 (0.1%) |
| | PB | 27219 (17.5%) | 1822 (6.7%) |
| | PB-like | 19 (0%) | 0 (0%) |
| | RMB | 32321 (20.7%) | 98 (0.3%) |

#### Extended Data Table 6. Description of monoclonal antibodies derived from GC B cells and LNPCs

| | | 01a (N=145) N (%) | 02a (N=410) N (%) | 04 (N=124) N (%) | 07 (N=201) N (%) | 10 (N=249) N (%) | 13 (N=108) N (%) | 20 (N=99) N (%) | 22 (N=167) N (%) |
|---|---|---|---|---|---|---|---|---|---|
| Heavy chain isotype | IGHG | 109 (75.2%) | 387 (94.4%) | 102 (82.3%) | 146 (72.6%) | 200 (80.3%) | 102 (94.4%) | 90 (90.9%) | 144 (86.2%) |
| | IGHA | 35 (24.1%) | 11 (2.7%) | 15 (12.1%) | 54 (26.9%) | 43 (17.3%) | 5 (4.6%) | 6 (6.1%) | 22 (13.2%) |
| | IGHM | 1 (0.7%) | 11 (2.7%) | 7 (5.6%) | 1 (0.5%) | 2 (0.8%) | 1 (0.9%) | 3 (3%) | 1 (0.6%) |
| | IGHD | 0 (0%) | 0 (0%) | 0 (0%) | 0 (0%) | 0 (0%) | 0 (0%) | 0 (0%) | 0 (0%) |
| Light chain isotype | IGKC | 102 (70.3%) | 223 (54.4%) | 61 (49.2%) | 133 (66.2%) | 183 (73.5%) | 66 (61.1%) | 69 (69.7%) | 73 (43.7%) |
| | IGLC | 43 (29.7%) | 187 (45.6%) | 63 (50.8%) | 68 (33.8%) | 66 (26.5%) | 42 (38.9%) | 30 (30.3%) | 94 (56.3%) |
| Compartment | GC B cell | 85 (58.6%) | 248 (60.5%) | 95 (76.6%) | 134 (66.7%) | 151 (60.6%) | 68 (63%) | 59 (59.6%) | 82 (49.1%) |
| | LNPC | 60 (41.4%) | 162 (39.5%) | 29 (23.4%) | 67 (33.3%) | 98 (39.4%) | 40 (37%) | 40 (40.4%) | 85 (50.9%) |
| Time point | Week 4 | | | 45 (36.3%) | | | | | |
| | Week 5 | | 180 (43.9%) | | | | | | |
| | Week 7 | 84 (57.9%) | 230 (56.1%) | 79 (63.7%) | 142 (70.6%) | 115 (46.2%) | 67 (62%) | 34 (34.3%) | 108 (64.7%) |
| | Week 15 | 61 (42.1%) | | | 59 (29.4%) | 134 (53.8%) | 41 (38%) | 65 (65.7%) | 59 (35.3%) |

### Extended Data Table 7. Processing of BCR and 5' gene expression data from scRNA-seq

| Participant | Time point | Compartment | BCR | | 5' gene expression | | | |
|---|---|---|---|---|---|---|---|---|
| | | | Pre QC number of cells | Post QC number of cells | Pre-QC number of cells | Post-QC number of cells | Median number of UMIs per cell | Median number of genes per cell |
| 01a | Week 4 | PB | 6211 | 5562 | 8106 | 6251 | 14020 | 2470 |
| | Week 7 | FNA | 5637 | 5198 | 16493 | 15065 | 10069 | 3904 |
| | Week 15 | FNA | 5044 | 4667 | 14544 | 13664 | 9803 | 3908 |
| 02a | Week 4 | PB | 3981 | 3016 | 2847 | 1952 | 8589.5 | 1967.5 |
| | Week 5 | FNA | 18943 | 16664 | 27065 | 24827 | 22731 | 8519 |
| | Week 7 | FNA | 26710 | 20982 | 39293 | 35052 | 20904 | 7936 |
| 04 | Week 4 | PB | 5347 | 4321 | 8571 | 4486 | 12855.5 | 2546 |
| | Week 4 | FNA | 5205 | 4868 | 11272 | 10312 | 11384 | 4267 |
| | Week 7 | FNA | 5771 | 5142 | 13993 | 13063 | 10733 | 3926 |
| 07 | Week 4 | PB | 4123 | 3610 | 6284 | 3745 | 14059 | 2602 |
| | Week 7 | FNA | 7284 | 6515 | 14466 | 13282 | 10986 | 4173 |
| | Week 15 | FNA | 8343 | 7457 | 16367 | 15340 | 10717 | 4183 |
| 10 | Week 4 | PB | 3228 | 2535 | 2336 | 1565 | 13046 | 2386 |
| | Week 7 | FNA | 20154 | 18088 | 33178 | 30545 | 22459.5 | 8451 |
| | Week 15 | FNA | 23419 | 20133 | 35386 | 31771 | 23104 | 8657.5 |
| 13 | Week 4 | PB | 2241 | 1897 | 3072 | 2258 | 12423.5 | 2433 |
| | Week 7 | FNA | 8448 | 7320 | 14126 | 13264 | 10978 | 4145 |
| | Week 15 | FNA | 3098 | 2716 | 8730 | 6927 | 5615 | 2186 |
| 20 | Week 4 | PB | 2962 | 2175 | 2011 | 1346 | 13254 | 2343 |
| | Week 7 | FNA | 5086 | 4595 | 15171 | 14459 | 10989.5 | 3758 |
| | Week 15 | FNA | 8754 | 7739 | 18876 | 17299 | 11520.5 | 4065 |
| 22 | Week 4 | PB | 6517 | 5313 | 9504 | 6850 | 11730 | 2251.5 |
| | Week 7 | FNA | 9580 | 7904 | 22879 | 21692 | 9700 | 3756 |
| | Week 15 | FNA | 4780 | 4138 | 16233 | 13141 | 10014.5 | 3802 |

**Extended Data Table 8. Processing of BCR reads from bulk-seq**

| Participant | Time point | Compartment | Cell count | Sequence count | | | |
|---|---|---|---|---|---|---|---|
| | | | | Input | Preprocessed | Post QC | Unique VDJ |
| 01a | Week 7 | GC B cell | 16920 | 1234529 | 4950 | 4018 | 2510 |
| | | LNPC | 3307 | 1350772 | 11799 | 9851 | 3592 |
| | Week 15 | GC B cell | 10440 | 1031639 | 1274 | 897 | 612 |
| | | LNPC | 2139 | 1659983 | 6337 | 5348 | 2122 |
| | Week 29 | BMPC | 16000000 | 2224729 | 142929 | 127601 | 65389 |
| 02a | Week 5 | GC B cell | 75639 | 2323018 | 60305 | 50108 | 28663 |
| | | LNPC | 7983 | 2062756 | 80227 | 61617 | 14236 |
| | Week 7 | GC B cell | 65172 | 2264893 | 47141 | 38634 | 21203 |
| | | LNPC | 8730 | 2041016 | 71862 | 54796 | 13202 |
| | Week 29 | BMPC | 200000 | 1132888 | 63177 | 55882 | 35438 |
| 04 | Week 4 | GC B cell | 27014 | 1191023 | 3115 | 2207 | 1779 |
| | | LNPC | 2312 | 1012560 | 9807 | 8276 | 2955 |
| | Week 7 | GC B cell | 37948 | 1354770 | 16773 | 13798 | 8757 |
| | | LNPC | 2495 | 917541 | 11273 | 9403 | 3361 |
| | Week 29 | BMPC | 200000 | 1526703 | 11664 | 10357 | 9241 |
| 07 | Week 7 | GC B cell | 67030 | 1330762 | 37376 | 32737 | 18798 |
| | | LNPC | 7131 | 1360131 | 29602 | 21560 | 6666 |
| | Week 15 | GC B cell | 36501 | 1160090 | 6994 | 5377 | 3762 |
| | | LNPC | 2108 | 1798209 | 19416 | 14404 | 3801 |
| | Week 29 | BMPC | 100000 | 1212296 | 19112 | 17647 | 14378 |
| 10 | Week 7 | GC B cell | 37570 | 2771591 | 43545 | 36356 | 19228 |
| | | LNPC | 13131 | 2747041 | 119842 | 90279 | 20947 |
| | Week 15 | GC B cell | 103688 | 2599601 | 78901 | 65390 | 34232 |
| | | LNPC | 15757 | 2472177 | 102827 | 77796 | 20465 |
| 13 | Week 7 | GC B cell | 26603 | 1448419 | 13749 | 11614 | 6413 |
| | | LNPC | 6798 | 1528468 | 35222 | 25799 | 6997 |
| | Week 15 | GC B cell | 38318 | 1424118 | 13795 | 11862 | 7757 |
| | | LNPC | 3801 | 1263065 | 26314 | 19482 | 5107 |
| | Week 29 | BMPC | 100000 | 3206444 | 21006 | 19123 | 14894 |
| 20 | Week 7 | GC B cell | 16773 | 1249336 | 10208 | 8084 | 4593 |
| | | LNPC | 3967 | 1248043 | 38803 | 28898 | 7185 |
| | Week 15 | GC B cell | 63961 | 999961 | 24525 | 20745 | 12230 |
| | | LNPC | 6877 | 1324665 | 3196 | 1998 | 1323 |
| | Week 29 | BMPC | 200000 | 1307413 | 602 | 491 | 380 |
| 22 | Week 7 | GC B cell | 17676 | 1344080 | 15060 | 12556 | 5925 |
| | | LNPC | 4823 | 1068783 | 26503 | 20779 | 4907 |
| | Week 15 | GC B cell | 48264 | 891196 | 22310 | 17775 | 8612 |
| | | LNPC | 10443 | 1290422 | 28342 | 19770 | 6665 |
| | Week 29 | BMPC | 200000 | 2185196 | 115201 | 105375 | 64106 |
